# Top Model Decision Tree: Selecting Segmentation Models for Reliable Quantitative Analysis in Low- and Ultralow-Dose CryoEM

**DOI:** 10.64898/2026.06.05.730486

**Authors:** Lynnicia N. Massenburg, Sita S. Madugula, Spenser R. Brown, Amber N. Bible, Chanda R. Harris, Lance X. Zhang, Kiara Parker, Scott T. Retterer, Jennifer L. Morrell-Falvey, Rama K. Vasudevan, Alexis N. Williams

**Author notes:** This manuscript has been authored by UT-Battelle, LLC, under contract DE-AC05-00OR22725 with the US Department of Energy (DOE). The United States Government retains and the publisher, by accepting the article for publication, acknowledges that the United States Government retains a nonexclusive, paid-up, irrevocable, worldwide license to publish or reproduce the published form of this manuscript, or allow others to do so, for the United States Government purposes. The Department of Energy will provide public access to these results of federally sponsored research in accordance with the DOE Public Access Plan(http://energy.gov/downloads/doe-public-access-plan).

## Abstract

Deep learning neural networks provide a powerful approach for segmenting low-contrast cryogenic electron microscopy (cryoEM) images. However, model performance can vary significantly across imaging conditions and may hinder downstream quantitative analyses. Here, we present a structured evaluation workflow to systematically screen segmentation models based on performance, inference speed, robustness across imaging conditions, and reliability of downstream quantitative measurements. Using the Bacterial Cell Envelope Thickness Tool (BCET) as a test case, we evaluate multiple architectures (YOLOv11, YOLO26, U-Net, Detectron2, and SAM3) under low-dose and ultralow-dose cryoEM conditions. While several models achieve high metrics, model choice strongly influences downstream measurements of envelope thickness. Models optimized for high F1-scores may produce unreliable segmentation masks from object crowding, interpolation artifacts or imaging conditions. Our results reveal distinct trade-offs between performance, speed, and robustness amongst models. YOLOv11 provides the highest fidelity membrane segmentation for quantitative measurements and the Meta-based model SAM3 offers improved robustness under ultralow-dose conditions with competitive inference performance. This work provides practical guidance for model selection in cryoEM workflows, emphasizing that optimal choice depends on experimental priorities and downstream analysis requirements rather than metrics alone. These findings are broadly relevant to cryoEM workflows as AI-based analysis expands beyond the biological sciences.

**Graphical Abstract:** 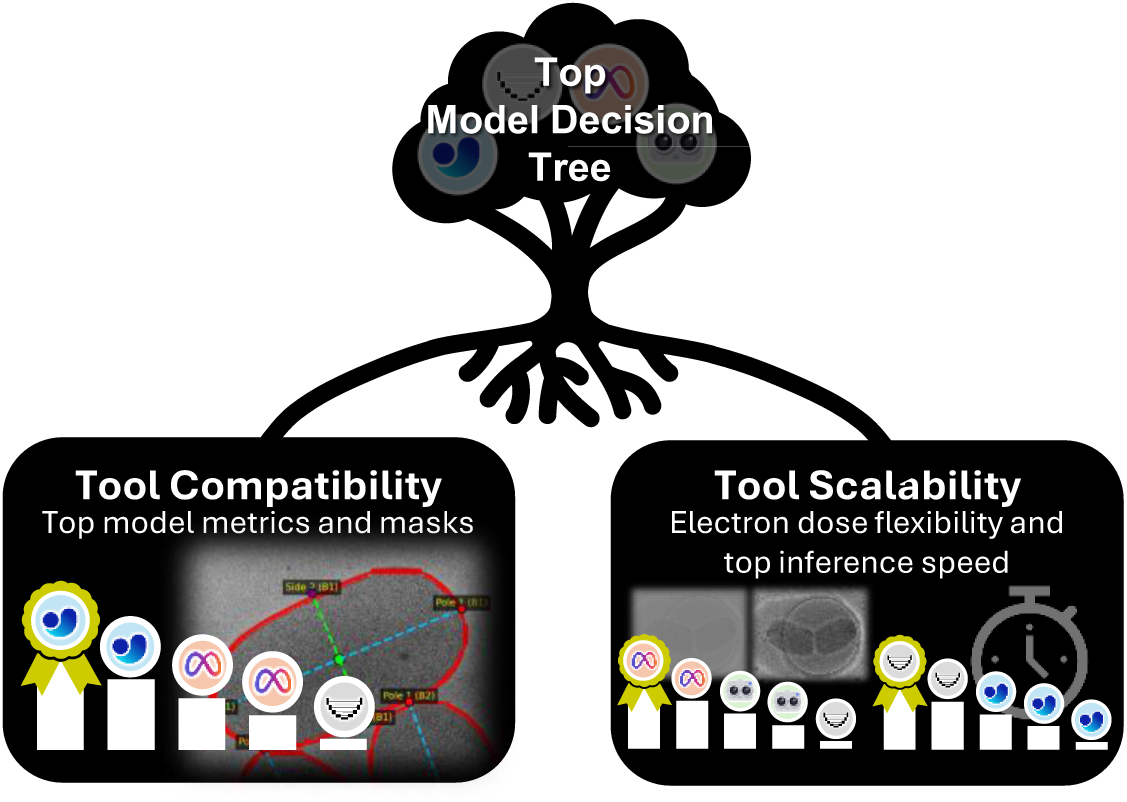

## Introduction

Biological feature segmentation in cryogenic electron microscopy (cryoEM) images represents a balance between achieving high-resolution imaging and minimizing electron beam damage to preserve sample integrity (Egerton, 2020). As a result, data collection is performed under low-dose conditions that produce low-contrast images and creates a strong need for deep learning-based computer vision methods to identify small or thin features of interest. We recently demonstrated success using a YOLOv11 architecture to identify bacterial inner and outer membranes and flagella under specific low-dose imaging conditions. Moreover, we developed the Bacterial Cell Envelope Thickness (BCET) Tool to quantitatively measure the distance between bacterial inner and outer membrane edges (e.g., the bacterial cell envelope thickness) using high-detail segmentation masks (Madugula et al., 2026). Importantly, this tool enables the evaluation of how segmentation model choice directly impacts quantitative measurements derived from low-dose cryoEM data.

Currently, no standardized model-manager framework exists to screen a collection of segmentation models in a structured environment. Instead, most studies adopt a one-model approach, tailoring a specific architecture to low-contrast biological images. As such, U-Net and related architectures such as Cellpose, have become canonical in biological segmentation tasks ranging from protein particle analysis to cell ultrastructure (Baumgartner et al., 2021; Bepler et al., 2020; Buchholz et al., 2019; Ronneberger et al., 2015; Sanchez-Garcia et al., 2021; Stringer et al., 2021). However, single-model approaches often struggle when applied to cell types or imaging conditions that differ from those represented in the training data, resulting in degraded performance (He et al., 2023). A one-for-many model called the Segment Anything Model (SAM) was developed to address this limitation with a “zero-shot” approach requiring no additional images with generalist model segmentation (Kirillov et al., 2023). This approach laid the foundation for Segment Anything for Microscopy (µSAM), CellSAM, SAM3-Adapter and Medical SAM3 to revamp the pre-trained SAM model on biological microscopy images for “zero-shot” inferencing of cell ultrastructures in optical and electron microscopy (Archit et al., 2025; T. Chen et al., 2025; Israel et al., 2025; Jiang et al., 2026). Nevertheless, these models still face challenges in resolving fine structural boundaries required for precise quantitative measurements in noisy cryoEM images, especially across different cell types and imaging conditions.

A critical limitation of current approaches is that segmentation models are typically evaluated using aggregate computational metrics such as F1-score, mAP50, IoU or Dice coefficient (Bishop, 2007; Everingham et al., 2010; Lin et al., 2014; Manning et al., 2008; Shelhamer et al., 2017; Sudre et al., 2017; Taghanaki et al., 2019; Travieso et al., 2024; Zou et al., 2004). However, these metrics quantify agreement with annotated ground truth and do not capture how segmentation errors propagate into downstream quantitative measurements. In workflows where segmentation outputs are used to derive physical parameters (e.g., cell envelope thickness, morphology, or defect characterization) even small deviations in mask geometry can lead to substantial measurement inaccuracies. As a result, models with similar performance metrics may produce significantly different scientific outcomes from the masks. This disconnect highlights the need for evaluation strategies that account for both segmentation performance and measurement reliability.

To address these challenges, we introduce a task-aware model selection framework that explicitly links segmentation outputs to downstream quantitative measurement fidelity. Using the BCET tool as a test case, we systematically evaluate multiple segmentation architectures (YOLOv11, YOLO26, U-Net, Detectron2, and SAM3) under consistent low-dose and ultralow-dose cryoEM conditions. Rather than identifying a single best-performing model based on aggregate metrics, this framework characterizes trade-offs between segmentation accuracy, inference speed, robustness across imaging conditions, and the reliability of derived measurements. We use a stepwise approach to select imaging modes, mask evaluation and metrics evaluation criteria to select the top model based on user goals (Fig. 1). Our results demonstrate a consistent decoupling between standard segmentation metrics and measurement accuracy, emphasizing that optimal model selection must be guided by application-specific requirements rather than metric performance alone. While demonstrated for bacterial membrane segmentation, this approach is broadly applicable to microscopy workflows in both biological and materials systems where segmentation serves as an intermediate step for quantitative analysis.

**Fig. 1.**
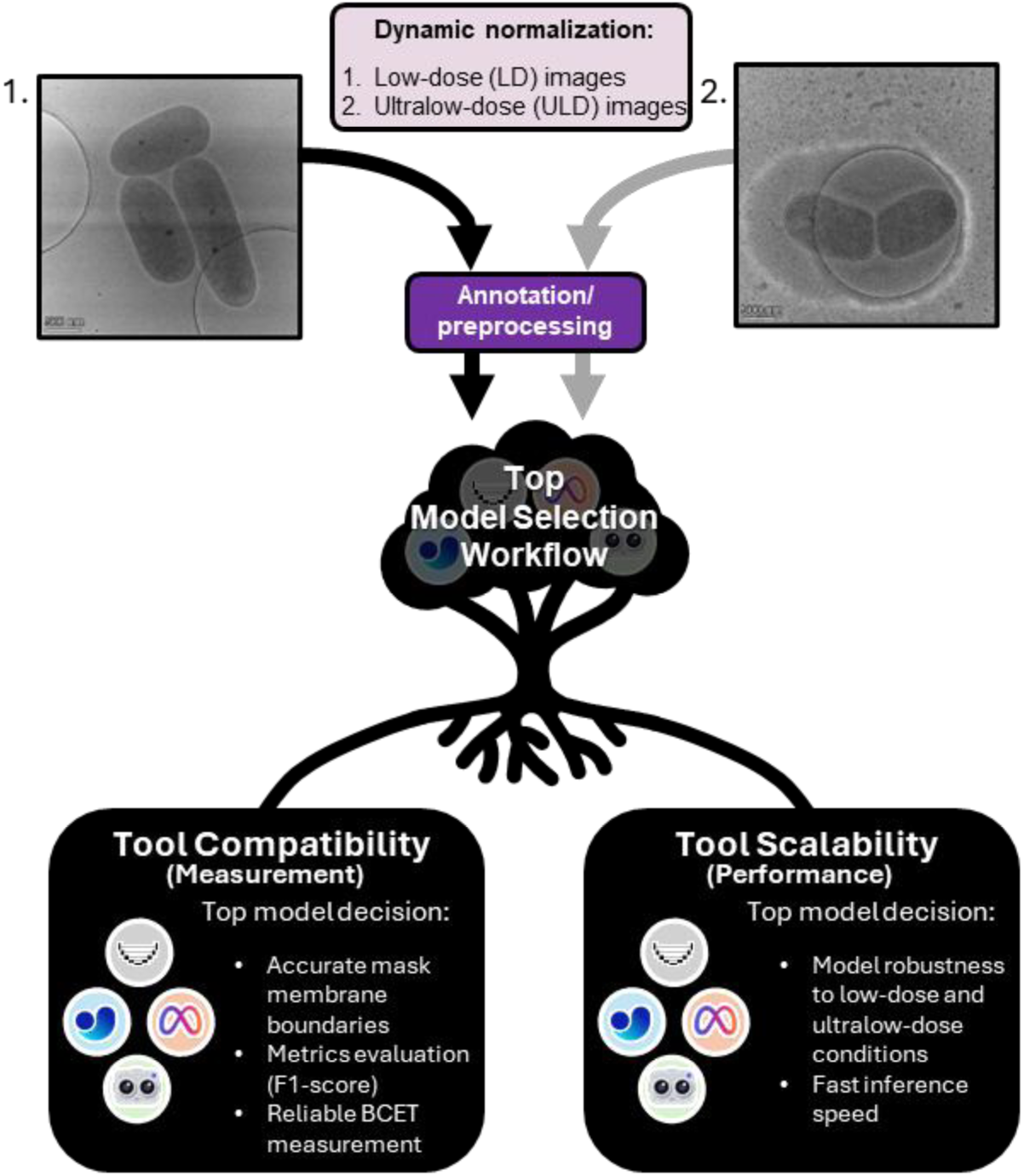
Schematic of the Top Model Decision Tree workflow. Low-dose (LD) and ultralow-dose (ULD) cryoEM images were processed through a common pipeline to generate segmentation masks and evaluation metrics. Models were then compared based on both quantitative performance metrics and qualitative mask quality to guide model selection for downstream applications. This workflow enables selection of models optimized for measurement reliability (tool compatibility) and inference speed and robustness (tool scalability). Scale bar is 500 nm.

## Materials and Methods

### Bacterial Growth

*Pantoea* sp. YR343 wild type (WT) used in this study was a Gram negative soil microbe that was isolated from the rhizosphere of native *Populus deltoides* trees in North Carolina (Bible et al., 2016). Single colonies of *Pantoea* sp. YR 343 were cultured and prepared for low-dose imaging as described in Madugula et al.(Madugula et al., 2026). Additional colonies of *Pantoea* sp. YR 343 were cultured in MOPS + succinate at 29 °C. Cells were collected and analyzed in stationary phase after overnight growth (24 hr).

### CryoEM Sample Preparation

Samples used for low-dose imaging were described in Madugula et al. (Madugula et al., 2026). For the ultralow-dose imaging, 10 μL aliquots of overnight bacterial samples grown in MOPS + succinate minimal media were applied to glow-discharged Quantifoil® 2/2 gold grids, blotted with a VitroBot Mark IV (Thermo Fisher Scientific, blot force −4 for 4 seconds at 100% humidity and 22 °C), and vitrified by plunge-freezing in liquid ethane/propane cooled by liquid nitrogen.

### CryoEM Data Collection

Manual cryoEM image collection was described in Madugula et al. for low-dose images collected at 40 e⁻/Å^2^ at –5 μm defocus using the Falcon 3EC direct electron detector (Thermo Fisher Scientific, counting mode) on the Thermo Fisher Scientific Krios G4 operated in nanoprobe TEM mode (Madugula et al., 2026). We used automated cryoEM imaging at an ultralow-dose rate to demonstrate unsupervised high-speed imaging. The Thermo Fisher Scientific Krios G4 was operated in nanoprobe TEM mode with a 70 μm C2 aperture under cryogenic conditions. A set of bacterial samples were imaged using an ultra-low dose at 6 e⁻/Å^2^ and –10 μm defocus with the Falcon 3EC direct electron detector (Thermo Fisher Scientific, integrated mode). The defocus value corresponds to the underfocus of the objective lens to enhance image contrast. We increased ultralow-dose image contrast by changing defocus from –5 μm (used in low-dose imaging) to –10 μm to sharpen membrane fine boundaries. Low magnification atlas images of the 3 mm grid were collected using the Thermo Fisher EPU software. Grid squares of interest were selected in EPU with an automated array imaging routine. Automated image arrays that were set to image empty carbon were removed from the imaging routine. Images were collected at 8700x magnification (0.8986 nm/pix) to identify bacterial features of interest. Data were automatically collected for bacteria in minimal MOPS + succinate media at stationary phase for ultra-low dose images (216 micrographs). A total of 122 images were used after discarding images with grid cracks or thick ice contamination.

### Pre-processing and Data Annotation

Preprocessing and data annotation on 149 low-dose images were described previously in Madugula et al (Madugula et al., 2026) with the addition of inter-linear downsizing to 1024 × 1024 and a dataset export to COCO format. Ultralow-dose cryoEM 4096 × 4096 images were dynamically contrast-normalized (adaptive-normalization) with variance-based percentile scaling and output as lossless 8-bit PNGs to preserve high-detail cryoEM information. These images were then loaded into Roboflow for manual annotation (Ciaglia et al., 2022). The inner membrane (IM) and outer membrane (OM) were annotated as filled polygon contours. A total of 122 images were split into 80% training, 5% validation, and 15% test sets. Datasets were inter-linear downsized to 640 × 640 or 1024 × 1024 and training images were augmented as described in Table 1 and 3x multiplied to address limited image accessibility. Datasets were exported in YOLOv11 and YOLO26 format as YOLO txt/polygons as well as exported in COCO format for JSON annotations used with U-Net, Detectron2 and SAM3. Only train and valid images were used for fine-tuning pre-trained base models. Instance segmentation models (YOLOv11, YOLO26, Detectron2 and SAM3) and semantic segmentation models (U-Net) were generated with both sets of downsized images as summarized in Table 1. Low-dose and ultralow-dose images were separately used in model generation to generate low-dose only (LD) model sets or ultralow-dose only (ULD) model sets. The shorthand LD and ULD will only be used in reference to the model set seeds for clarity. Each LD and ULD model set (or seed) had two additional model subsets based on training image size: one model subset at 640 × 640 and another subset at 1024 × 1024. Both subsets were used to determine the top models in their respective image condition seed. All image downsampling and upsampling in model training underwent inter-linear (also called bilinear) interpolation. Google Colab Pro with GPU (Pytorch backend) was used for all model training.

**Table 1.**
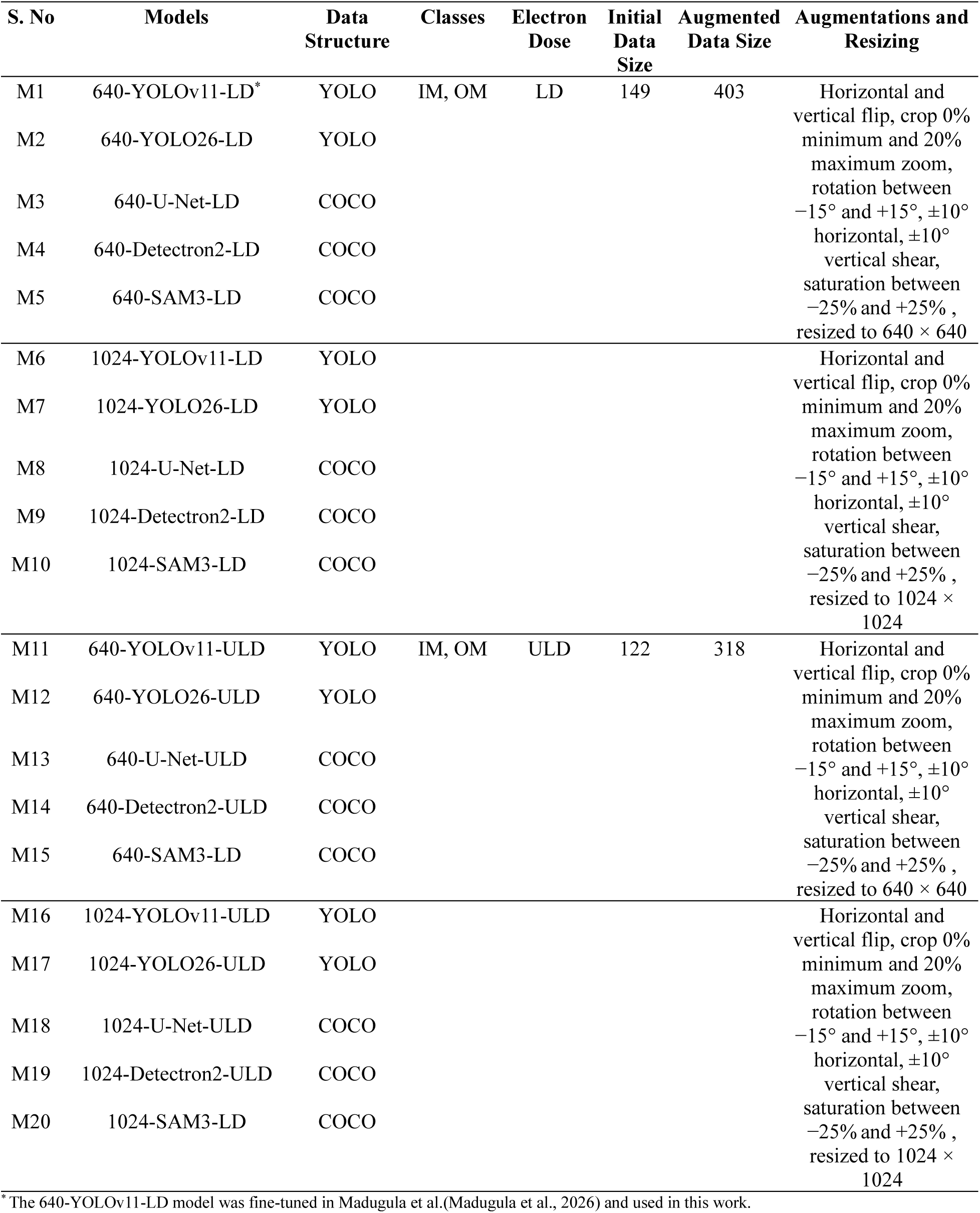
Summary of datasets, augmentations, and resizing applied for model building the outer membrane and inner membrane for bacterial cell envelope thickness.

| S. No | Models | Data Structure | Classes | Electron Dose | Initial Data Size | Augmented Data Size | Augmentations and Resizing |
| --- | --- | --- | --- | --- | --- | --- | --- |
| M1 | 640-YOLOv11-LD* | YOLO | IM, OM | LD | 149 | 403 | Horizontal and vertical flip, crop 0% minimum and 20% maximum zoom, rotation between $-15^\circ$ and $+15^\circ$, $\pm 10^\circ$ horizontal, $\pm 10^\circ$ vertical shear, saturation between $-25\%$ and $+25\%$, resized to $640 \times 640$ |
| M2 | 640-YOLO26-LD | YOLO |  |  |  |  |  |
| M3 | 640-U-Net-LD | COCO |  |  |  |  |  |
| M4 | 640-Detectron2-LD | COCO |  |  |  |  |  |
| M5 | 640-SAM3-LD | COCO |  |  |  |  |  |
| M6 | 1024-YOLOv11-LD | YOLO | | | | | Horizontal and vertical flip, crop 0% minimum and 20% maximum zoom, rotation between $-15^\circ$ and $+15^\circ$, $\pm 10^\circ$ horizontal, $\pm 10^\circ$ vertical shear, saturation between $-25\%$ and $+25\%$, resized to $1024 \times 1024$ |
| M7 | 1024-YOLO26-LD | YOLO |  |  |  |  |  |
| M8 | 1024-U-Net-LD | COCO |  |  |  |  |  |
| M9 | 1024-Detectron2-LD | COCO |  |  |  |  |  |
| M10 | 1024-SAM3-LD | COCO |  |  |  |  |  |
| M11 | 640-YOLOv11-ULD | YOLO | IM, OM | ULD | 122 | 318 | Horizontal and vertical flip, crop 0% minimum and 20% maximum zoom, rotation between $-15^\circ$ and $+15^\circ$, $\pm 10^\circ$ horizontal, $\pm 10^\circ$ vertical shear, saturation between $-25\%$ and $+25\%$, resized to $640 \times 640$ |
| M12 | 640-YOLO26-ULD | YOLO |  |  |  |  |  |
| M13 | 640-U-Net-ULD | COCO |  |  |  |  |  |
| M14 | 640-Detectron2-ULD | COCO |  |  |  |  |  |
| M15 | 640-SAM3-LD | COCO |  |  |  |  |  |
| M16 | 1024-YOLOv11-ULD | YOLO | | | | | Horizontal and vertical flip, crop 0% minimum and 20% maximum zoom, rotation between $-15^\circ$ and $+15^\circ$, $\pm 10^\circ$ horizontal, $\pm 10^\circ$ vertical shear, saturation between $-25\%$ and $+25\%$, resized to $1024 \times 1024$ |
| M17 | 1024-YOLO26-ULD | YOLO |  |  |  |  |  |
| M18 | 1024-U-Net-ULD | COCO |  |  |  |  |  |
| M19 | 1024-Detectron2-ULD | COCO |  |  |  |  |  |
| M20 | 1024-SAM3-LD | COCO |  |  |  |  |  |

\*The 640-YOLOv11-LD model was fine-tuned in Madugula et al.(Madugula et al., 2026) and used in this work.

### YOLOv11 Model Training and Hyperparameters

YOLOv11n-seg (114 layers, 2,834,958 parameters, 0 gradients, 9.6 GFLOPs) (Khanam & Hussain, 2024) was used as the starting pre-trained object-level instance segmentation model for fine-tuning implemented using the Ultralytics library (Jocher & Qiu, 2024). Batch normalization was bypassed as the images were adaptive-normalized during pre-processing. Models were trained at 100 epochs at a 16-batch size, learning rate of 0.001, IoU threshold of 0.5 and a cosine learning rate schedule (lrf) of 0.01 with an AdamW optimizer while the remaining parameters are kept to default. Loss functions tracked box loss, classification loss, and distribution focal loss (DFL). The best models were saved as 640-YOLOv11-LD, 1024-YOLOv11-LD, 640-YOLOv11-ULD, and 1024-YOLOv11-ULD.

### YOLO26 Model Training and Hyperparameters

The image segmentation training routine was similar to YOLOv11 apart from using YOLO26n-seg (139 layers, 2,689,274 parameters, 0 gradients, 9.0 GFLOPs)(Sapkota et al., 2025) as the starting pre-trained instance segmentation model for fine-tuning within the Ultralytics library. The best models with the highest mAP50-95 were saved as 640-YOLO26-LD, 1024-YOLO26-LD, 640-YOLO26-ULD, and 1024-YOLO26-ULD.

### U-Net Model Training and Hyperparameters

A U-Net-based convolutional neural network (CNN) was implemented with a five-level encoder–decoder architecture. The encoder was comprised of repeated blocks of two 3 × 3 convolutions followed by Rectified Linear Unit (ReLU) activation, with feature depths increasing from 32 to 512 channels (32, 64, 128, 256, 512). Within the encoder, dropout (p = 0.1) was applied to mitigate overfitting, and downsampling is performed via 2 × 2 max pooling, reaching a bottleneck layer with 1024 channels. The decoder restored spatial resolution through bilinear upsampling and skip connections to the corresponding encoder layers, followed by convolutional blocks mirroring the encoder structure (Ronneberger et al., 2015). Batch normalization omitted as the images were pre-processed using adaptive normalization. The final 1×1 convolutional layer produced two output channels corresponding to the IM and OM segmentation classes. To optimize both class-wise region overlaps and pixel-wise classification, a combined Dice-BCE loss function was employed. Early stopping is integrated with a patience of 10 epochs and a minimum delta of 0.001. Test images were trained for 180 epochs at a 4-batch size and a learning rate of 1 × 10^−4^ with the Adam optimizer. Early stopping was integrated at a patience of 10 epochs at a Δmin of 0.001. Loss functions included mask loss, IoU loss and Dice loss. The best models with the lowest Dice-BCE loss were selected and saved as 640-U-Net-LD, 1024 U-Net-LD, 640-U-Net-ULD, and 1024-U-Net-ULD.

### Detectron2 Model Training and Hyperparameters

Models were trained using a standard Detectron2 Mask R-CNN object-level instance segmentation algorithm (He et al., 2017). Training was conducted at 8,000 iterations with a batch size of 2 and a mask head resolution of 56. Batch normalization was omitted to maintain the intensity distributions established during the adaptive-normalization pre-processing stage. To maximize the two-class IM and OM membrane detection and minimize optimization oscillations, a custom warmup period was implemented for the first 200 iterations with a 0.4 warmup factor from a 2.5 × 10^−4^ learning rate to a final 1 × 10^−4^ learning rate (L. Chen et al., 2025). The model utilized a Stochastic Gradient Descent (SGD) optimizer with early stopping integrated at a patience of five validation rounds (conducted every 200 iteration periods per round) at a Δmin of 0.001. The total loss was calculated as the summation of classification loss, bounding box regression loss, segmentation mask loss, Region Proposal Network (RPN) objectness loss, and RPN box regression loss. The best models with the highest Average Precision (AP) were saved as 640-Detectron2-LD, 1024-Detectron2-LD, 640-Detectron2-ULD and 1024-Detectron2-ULD.

### SAM3 Model Training and Hyperparameters

The SAM3 (Meta AI) pre-trained models were fine-tuned using the object-level instance segmentation algorithm (Carion et al., 2025; Kirillov et al., 2023). Training was configured using a standard dataset descriptor (roboflow_v100.yaml) within the SAM3 repository that was customized to support mask-level segmentation. Batch normalization was omitted to maintain the intensity distributions in the pre-processing adaptive-normalization. To maintain the structural alignment required by the transformer-based image encoder, all images were resized to SAM’s native 1008 × 1008 resolution. Standard 1024 x 1024 dimensions were avoided to prevent positional embedding mismatches and training instability. The default pretrained Hugging Face Meta SAM3 checkpoint (facebook/sam3) using the SAM3 repository at commit f6e51f5. The base weights were initialized via the Hugging Face repository. Training was conducted for 20 epochs with a batch size of 4using the AdamW optimizer, with a vision backbone learning rate of 2.5 × 10^−5^ and a global learning rate of 5 × 10^−7^. To improve generalization, a vision learning rate decay of 0.9 and a weight decay of 0.1 were applied. The loss function was a weighted sum of bounding box regression, classification, and mask segmentation loss to optimize error balance. The final models were selected based on the lowest validation Dice loss. The best models with the lowest loss, mask loss and dice loss were selected and saved as 640-SAM3-LD, 1024-SAM3-LD, 640-SAM3-ULD, and 1024-SAM3-ULD.

### Model Evaluation

Model performance was evaluated based on precision, recall, and F1-score (F1) in addition to Average Precision at IoU thresholds of 0.5 (mAP50) for instance segmentation models as well as Mean IoU and Mean Dice for both semantic and instance segmentation models. The latter two metrics quantify the overlap between predicted and ground-truth segmentations while emphasizing different aspects of segmentation agreement, as shown in Supplementary Table 1 and Supplementary Table 2. For framework clarity, the F1-score was used as the top model metric in this work to compare semantic and instance segmentation models. Training was tracked using segmentation loss to optimize object shape accuracy. Valid images were resized to match the training dataset and used in the model evaluation for both low-dose and ultralow-dose images.

### Top Model Decision Tree Framework

The Top Model Decision Tree framework was built in Google Colab with A100 GPU enabled to store and inference a model collection and output a csv metrics table and a masks folder. Our model collection consists of two model sets called seeds: A low-dose (LD) model seed and an ultralow-dose (ULD) model seed designed as model repositories of all models used in this work (YOLOv11, YOLO26, U-Net, Detectron2, and SAM3) trained from either the low-dose train and valid images or ultralow-dose train and valid images (Supplementary Fig. 2). Low-dose and ultralow-dose test images were reserved for unbiased test metrics and masks used in the Top Model Decision Tree. Metrics were aggregated into a “top_model_tree.csv” table to compare model performance (Supplementary Fig, 2b). Test images were inter-linear resized to 640 × 640 or 1024 × 1024 to test for inference speed (640) or mask detail (1024). All predicted masks were inferenced at 1024 × 1024 image size at a 0.5 confidence threshold regardless of input model performance to maintain image detail. The masks were used in the Bacterial Cell Envelope Thickness (BCET) tool for quantitative measurements as previously described (Madugula et al., 2026). Predicted masks used in the BCET tool were also inferenced at 1024 × 1024 image size a 0.5 confidence threshold to maintain image detail. YOLOv11 and YOLO26 predicted OM masks were generated as a boundary ring layer instead of a filled mask that initially did not align with the ground truth OM filled mask. To compare ground truth OM mask to the predicted OM mask with test images, the ground truth boundary ring layer was created by subtracting the ground truth IM filled mask from the ground truth OM filled mask to generate the ground truth OM boundary layer mask. Overall best and per-image macro-averaged metrics were then calculated from ground truth and predicted IM filled masks and OM boundary layer masks for YOLOv11 and YOLO26. All other models predicted OM masks as filled masks. Model inference speed testing was performed on an NVIDIA A100-SXM4-40GB (1 GPU, CUDA v 12.8, computer capacity of 8.0) in a separate Google Colab at a 0.5 confidence threshold to track the average time per image and the frames per second (FPS) of low-dose and ultralow-dose test images at the two resize conditions mentioned above. Model speeds were tested in-tandem on the same GPU instance.

## Results and Discussion

### Top Model Decision Tree

We developed the Top Model Decision Tree to evaluate how model choice influences downstream quantitative measurements, rather than to identify a single best-performing model based on aggregate metrics (Fig. 2). We first evaluated model performance on low-dose cryoEM images collected at 40 e⁻/Å^2^. After applying dynamic normalization, we trained a low-dose (LD) model seed using fewer than 150 images and evaluated multiple architectures (YOLOv11, YOLO26, U-Net, Detectron2, and SAM3) under the identical 80:5:15 train, valid and test split. All models achieved strong validation performance (F1 = 0.898–0.997).

**Fig. 2.**
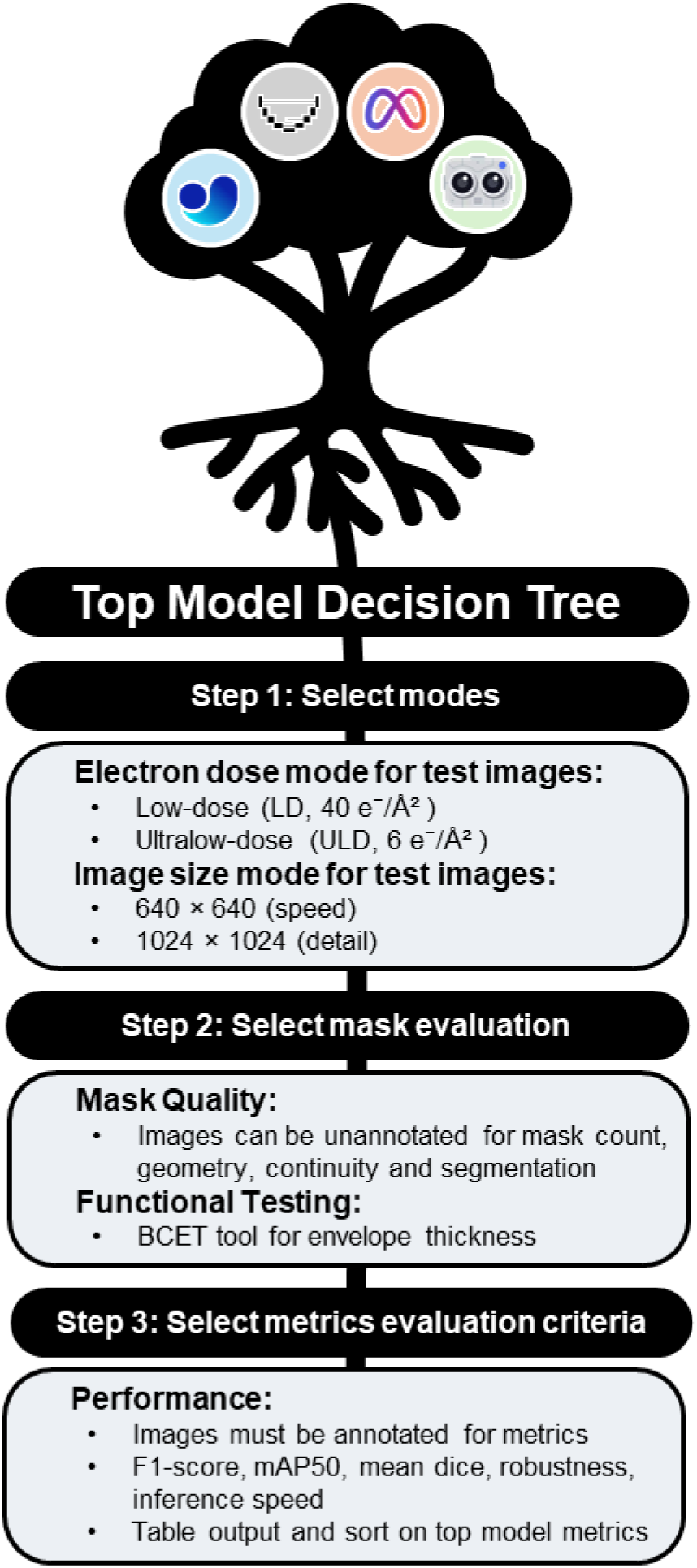
Top Model Decision Tree workflow. Overview of the model evaluation and selection pipeline. Pre-trained segmentation models and annotated cryoEM test images were processed to generate segmentation masks and evaluation metrics. Models were compared based on both quantitative metrics and qualitative mask characteristics to guide selection. Stepwise workflow for model selection. Users select a model seed (low-dose or ultralow-dose) and evaluate masks (.png file outputs) and metrics (.csv table outputs). The workflow outputs were used to assess model performance based on segmentation accuracy, inference speed, and mask quality relevant to downstream quantitative measurements.

We next introduced ultralow-dose (ULD) imaging at 6 e⁻/Å^2^ to evaluate robustness and scalability. ULD imaging reduced acquisition time from ~2 minutes per image (LD) to ~6 seconds per image, corresponding to a ~20× increase in throughput speed (Mendez et al., 2019). Despite reduced signal, dynamic normalization preserved membrane visibility, and models trained on ULD data achieved comparable validation performance (F1 = 0.883–1.0). These results show that reducing electron dose does not significantly compromise segmentation performance, enabling faster data acquisition without sacrificing model accuracy (Kazimi & Sandfeld, 2024). This finding supports the use of ultralow-dose imaging for high-throughput cryoEM workflows.

The workflow operates as a stepwise process (Fig. 2), generating segmentation masks and evaluating outputs using both quantitative metrics and qualitative mask inspection. We assessed each model’s ability to trace membrane boundaries, preserve object separation, and maintain consistent geometry, as these properties directly affect downstream quantitative measurements. These results show that segmentation performance appears similar across imaging conditions when evaluated using F1-scores (Supplementary Fig 3-8). However, models differ in inference speed and in their ability to accurately trace membrane boundaries for reliable quantitative measurements. These findings demonstrate that evaluating segmentation models based on performance metrics alone is insufficient and that model selection must account for downstream measurement requirements and imaging constraints.

### Tool Compatibility: Top Models for the Bacterial Cell Envelope Thickness (BCET) Tool

We evaluated model compatibility with the BCET tool using low-dose (LD) test images at 1024 × 1024 resolution. Based on F1-score, YOLOv11 models ranked highest, followed by SAM3 and U-Net (Fig. 3a, Table 2). We then used the BCET tool to evaluate the ability of each model to accurately measure a biological feature of interest. The BCET tool computes cell envelope thickness by tracing inner membrane (IM) and outer membrane (OM) boundaries from segmentation masks. Accurate measurements therefore require continuous membrane contours, proper separation between adjacent bacteria, and consistent boundary geometry. Disruptions in these properties directly affect thickness calculations. YOLOv11 and SAM3 produced accurate and consistent thickness measurements, whereas U-Net generated malformed outputs despite high F1-scores (Fig. 3b). Visual inspection of segmentation masks (Supplementary Fig. 9) showed that models differed in their ability to preserve membrane continuity and object separation, which determined whether masks could support reliable BCET analysis. We initially hypothesized that mask representation (boundary versus filled OM masks) would affect BCET performance, since the tool was developed using boundary-layer masks from YOLOv11. However, we found that mask geometry and structural consistency were more important than mask format, as demonstrated by the strong performance of SAM3.

**Fig. 3.**
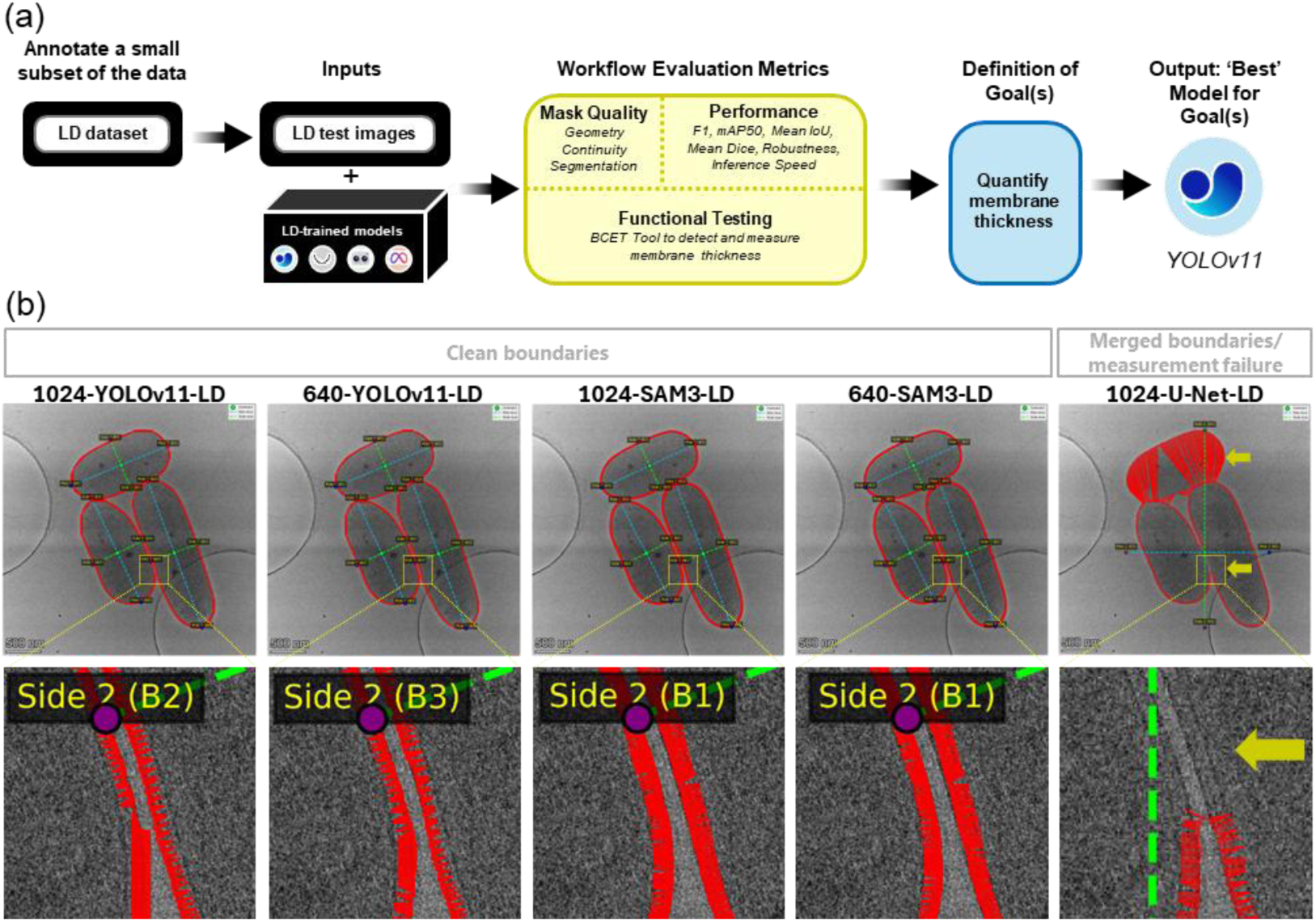
Compatibility of segmentation models with the Bacterial Cell Envelope Thickness (BCET) tool. (a) Workflow showing selection of the low-dose (LD) model seed and 1024 × 1024 test image size for evaluation within the Top Model Decision Tree. (b) Representative BCET outputs for top selected models, including segmentation masks and corresponding cell envelope thickness measurements. The yellow arrows correspond to merged boundaries that resulted in measurement failure. Despite similar F1-scores, models produce different mask geometries, resulting in differences in measurement accuracy. Scale bar is 500 nm.

**Table 2.** Quantitative analysis of the top five LD models in bacterial membrane segmentation of low-dose test images.

| S. No | Models | Mask<br>Consistency | BCET<br>Failure Rate | F1-Score** | mAP50 | Mean<br>IoU | Mean<br>Dice** |
| --- | --- | --- | --- | --- | --- | --- | --- |
| M6 | 1024-YOLOv11-LD | 20/20<br>(100%) | 2/20<br>(10%) | 0.999 | 0.995 | 0.877 | 0.930 |
| M1 | 640-YOLOv11-LD | 20/20<br>(100%) | 2/20<br>(10%) | 0.998 | 0.995 | 0.888 | 0.937 |
| M10 | 1024-SAM3-LD | 20/20<br>(100%) | 0/20<br>(0%) | 0.994 | 1.0 | 0.990 | 0.995 |
| M5 | 640-SAM3-LD | 20/20<br>(100%) | 0/20<br>(0%) | 0.994 | 1.0 | 0.989 | 0.994 |
| M8 | 1024-U-Net-LD* | 18/20<br>(90%) | 4/18<br>(22%) | 0.987 | - | 0.972 | 0.985 |

\*U-Net model evaluations do not generate mAP50 scores as the output is semantic segmentation, not instance segmentation for YOLOv11, YOLO26, Detectron2 and SAM3 models. Hence, the best F1-score was chosen as the top model metrics criteria.
\*\*F1-score represents the best F1-score across the confidence threshold, and the best Dice coefficient should equal the best F1-score. Mean Dice represents the average Dice coefficient with sensitivity to performance variations across a set of test images, and likewise the mean F1-score should equal the mean Dice coefficient. In addition, the best metrics were calculated across the dataset (global aggregation), whereas the mean metrics were calculated per image, and may cause a shift in the weighted scores that can lead to the mean value outperforming the best value.

We observed that image resizing contributed to substantial differences in model performance across architectures, particularly for U-Net. Bilinear interpolation introduced boundary blurring and overlapping artifacts in the 1024-U-Net model when downsampling images from 4096 × 4096 to 1024 × 1024, whereas inter-area interpolation improved boundary preservation (Supplementary Fig. 8). We did not implement the inter-area interpolation to maintain consistent preprocessing across models, ensuring that observed differences reflect model behavior rather than preprocessing variation (Long et al., 2015; Odena et al., 2016; Ronneberger et al., 2015; Sugawara et al., 2019). These interpolation-induced boundary artifacts become particularly apparent in images containing closely spaced bacteria. U-Net frequently merged adjacent cells at higher resolution (1024 × 1024), leading to inaccurate thickness measurements. Although higher resolution increased visual detail, it also led to boundary overprediction and loss of object separation. This behavior was associated with pixel-level (semantic) segmentation in U-Net, which does not explicitly model object instances and can result in merging of closely spaced bacteria. In contrast, reducing the input resolution to 640 × 640 improved object separation and produced more reliable BCET measurements, highlighting a trade-off between segmentation detail and quantitative measurement accuracy. Detectron2 generated consistent masks but exhibited edge oscillations that affected boundary precision, while YOLO26 showed sensitivity to image scaling and produced valid masks only at higher resolution. These results show that image resolution plays a critical role in segmentation performance. Higher-resolution inputs can degrade boundary fidelity and object separation, whereas lower-resolution inputs can improve consistency and support more reliable quantitative measurements (Sapkota & Karkee, 2025).

We next evaluated whether segmentation accuracy translated into consistent object detection and downstream quantitative measurements. Using a manually annotated set of 20 bacteria as ground truth, both YOLOv11 and SAM3 correctly identified all 20 objects, whereas lower-performing models exhibited under- or over-counting errors (Supplementary Fig. 10a and 10c; Supplementary Table 3). These discrepancies indicate that segmentation errors not only affect pixel-level accuracy but also propagate to object-level detection. To assess the impact of these differences on quantitative analysis, we compared radial cell envelope thickness profiles generated by the BCET tool across the same 20 bacteria (Fig. 4; Supplementary Fig. 10b and 10d). Models with accurate object detection produced consistent and biologically plausible thickness profiles, while models with segmentation errors showed increased variability and, in some cases, failed measurements (e.g., 0 nm thickness outputs). These failures were often associated with merged objects or disrupted membrane boundaries, which directly interfere with BCET analysis.

**Fig. 4.**
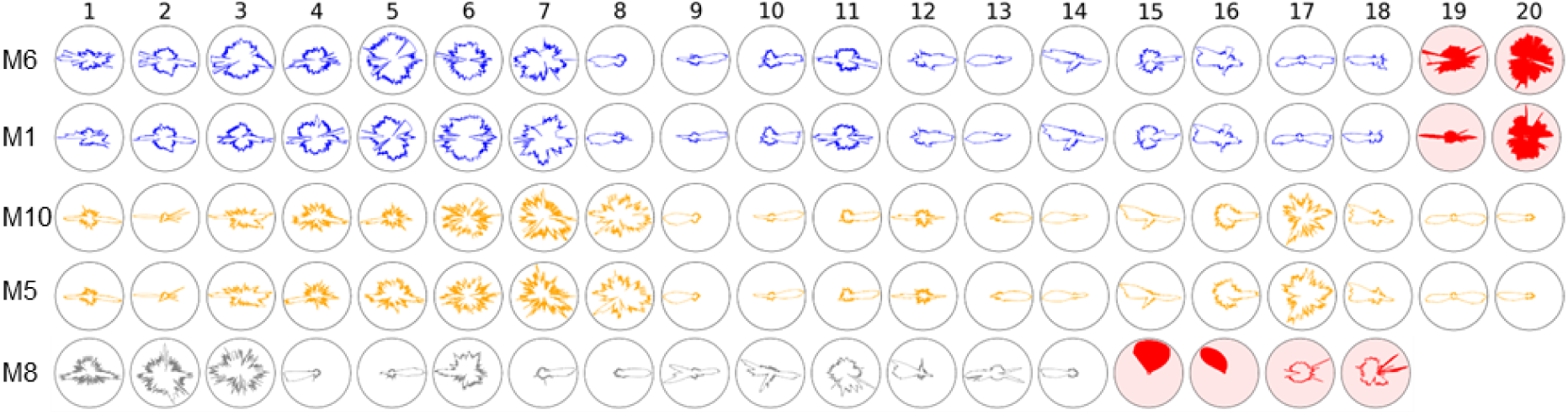
Model-dependent variability in quantitative cell envelope thickness measurements from low-dose cryoEM images. Radial cell envelope thickness profiles for 20 bacteria generated using the BCET tool from segmentation masks produced by 640-LD models. Models were ordered left-to-right based on successful thickness measurements (>0 nm, no merged bacterial neighbors). YOLOv11 (blue) and SAM3 (orange) produce consistent and biologically plausible thickness profiles across all bacteria, whereas other models exhibit increased variability and measurement failures. Radial plots corresponding to failed measurements (0 nm thickness or merged bacterial neighbors) were shown in red and were associated with segmentation errors such as merged objects or disrupted membrane boundaries. Plot colors indicate model architecture: YOLOv11 (blue), U-Net (gray) and SAM3 (orange). Plots were ranked top-down on best model (S. No. in Table 1) F1-score: (M6) 1024-YOLOv11-LD, (M1) 640-YOLOv11-LD, (M10) 1024-SAM3-LD, (M5) 640-SAM3-LD and (M8) 1024-U-Net-LD.

To quantitatively summarize these differences, we compared mask consistency and BCET tool measurement failure rates across the top five models (Table 2). Models that produced continuous boundaries and accurate object separation (YOLOv11 and SAM3) showed low variability and low measurement failures, whereas models with segmentation artifacts exhibited inconsistent masks and frequent invalid measurements (0 nm outputs and merged bacteria neighbors with U-Net). Together, these results demonstrated that accurate object detection is a prerequisite for reliable quantitative measurement. While counting-based approaches are commonly used in other microscopy modalities for population-level analysis (Halsted et al., 2022; Wang et al., 2021), the strength of this cryoEM-based workflow lies in linking object segmentation to structural measurements. By integrating segmentation metrics, mask evaluation, and BCET-derived outputs, we show that model choice directly impacts both object identification and measurement reliability, reinforcing the need for task-aware model selection.

### Tool Scalability: Accelerate Imaging and Inferencing Speed

Once we discovered that ULD imaging maintained segmentation performance in noisy data, we wanted to further evaluate how model choice affects robustness, inference speed, and segmentation consistency under these conditions. We first assessed how models trained on LD data generalize to ULD images. When we applied LD-trained models to ULD test images at 1024 × 1024 resolution, most models showed metric degradation and mask inconsistencies (Supplementary Fig. 11; Supplementary Table 3), demonstrating limited robustness to changes in imaging conditions. However, the LD-trained 640-SAM3 model retained relatively strong performance (F1 = 0.886) on ULD images, demonstrating greater robustness to imaging condition changes than other models (Fig 5a).

**Fig. 5.**
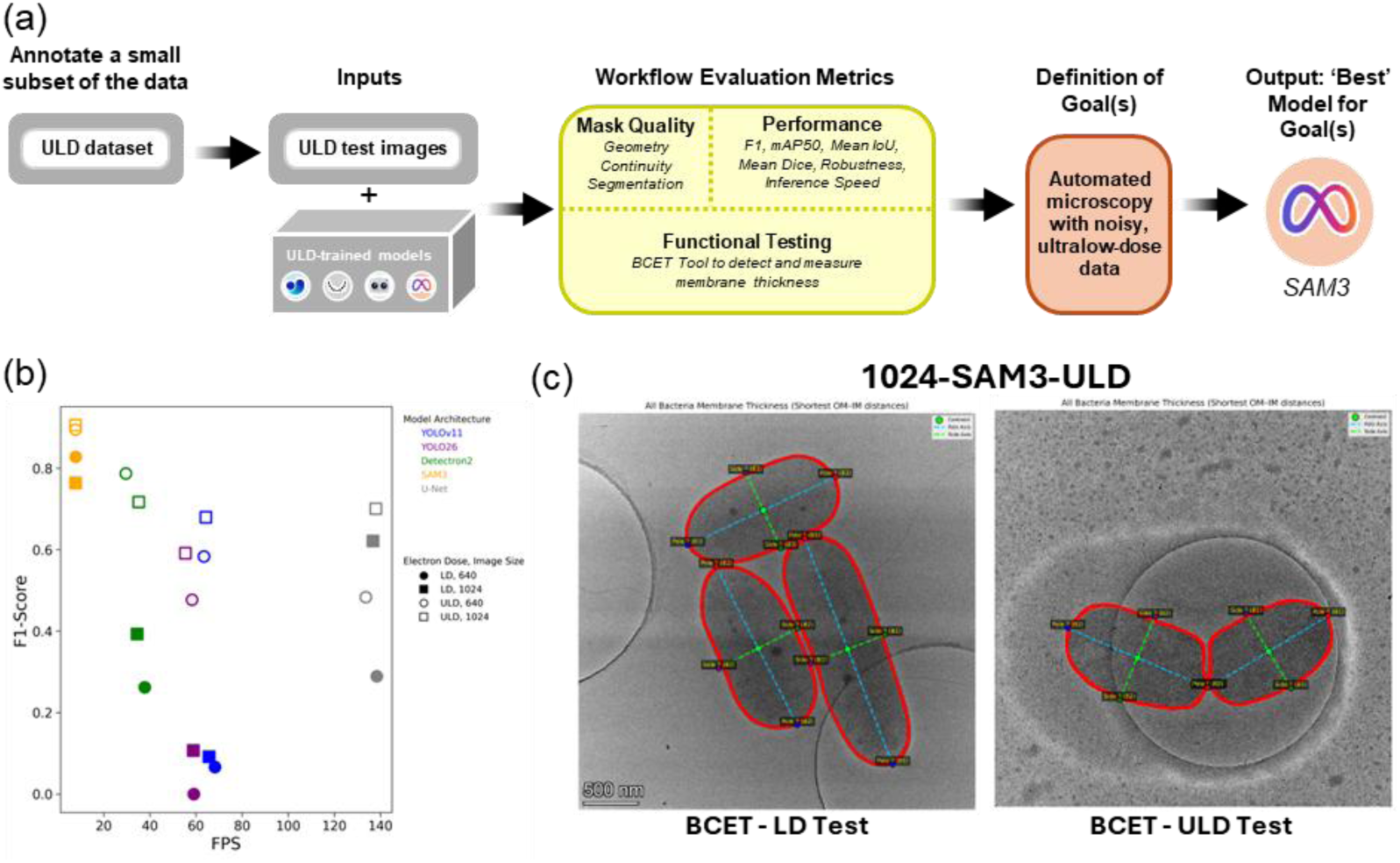
Scalability of segmentation and quantitative measurement under ultralow-dose (ULD) imaging conditions. (a) Workflow illustrating model selection based on segmentation accuracy, inference speed, and segmentation consistency. (b) Relationship between the best F1-score and inference speed (FPS) for ULD test images. (c) BCET outputs for the selected model (1024-SAM3-ULD) applied to low-dose (left) and ultralow-dose (right) images. Scale bar is 500 nm.

We next evaluated inference speed by reducing image size from 1024 × 1024 to 640 × 640. This reduction increased inference speed across all models. U-Net achieved ~140 FPS, while SAM3 remained slower at ~8 FPS (Fig. 5b; Supplementary Fig. 12). YOLOv11 and YOLO26 reached intermediate speeds (~60 FPS), while Detectron2 operated at ~30 FPS. These results highlight a clear trade-off between computational efficiency and segmentation performance.

Given the observed performance degradation of LD-trained models on ULD images, we evaluated whether training on ULD data improves model performance under these conditions. In the ULD trained instances SAM3 achieved the highest performance (F1 = 0.906 and 0.895 for 1024 and 640 models, respectively), outperforming LD-trained models. Detectron2 and U-Net followed, with lower F1-scores (0.787, 0.717, and 0.701). Although U-Net achieved the fastest inference speed, it showed reduced accuracy and poor segmentation consistency.

Model selection for scalability depended on three factors: segmentation accuracy (F1-score), inference speed, and segmentation consistency. Segmentation consistency emerged as a critical criterion, as models with high aggregate metrics (e.g., U-Net) produced unreliable outputs, with per-image mean F1-scores below 0.5 (Supplementary Table 4). Detectron2 provided a balanced trade-off between speed and consistency, whereas SAM3 delivered the highest accuracy and most consistent segmentation performance. This finding suggests that the improved SAM3 may outperform U-Net and Detectron2 in ultralow-dose conditions compared to a previous SAM build that underperformed in lower contrast medical images compared to U-Net (He et al., 2023).

Based on segmentation consistency and overall performance, we selected 1024-SAM3-ULD for BCET analysis across both low-dose and ultralow-dose images (Fig. 5c; Supplementary Fig 13-14, Supplementary Table 4). This model also performed strongly on low-dose data (F1 = 0.986), demonstrating robustness across imaging conditions. The selected model produced consistent BCET measurements and high-quality masks, outperforming other ULD models. More importantly, because ultralow-dose imaging operates at ~0.17 FPS (one frame every 6 seconds), SAM3 inference can be performed within acquisition time, enabling near–real-time analysis (Kuijper et al., 2015). This capability supports more efficient cryoEM workflows and enables integration of segmentation into live imaging pipelines for on-the-fly experiments and automated acquisition (Wang et al., 2025).

## Conclusions

The Top Model Decision Tree provides a structured approach for selecting segmentation models in cryoEM workflows based on application-specific requirements rather than aggregate performance metrics alone. By explicitly linking segmentation outputs to downstream quantitative analysis using the BCET tool, we demonstrate that models with similar F1-scores can produce substantially different measurement outcomes due to differences in mask geometry, object separation, and boundary fidelity. Our results highlight three key findings. First, segmentation performance metrics alone were insufficient for evaluating model suitability in scientific workflows since they do not capture errors that propagate into quantitative measurements. Second, model behavior varies systematically across architectures. YOLO-based models provide high-fidelity boundary representations for measurement tasks. U-Net offers fast inference with pixel-level confidence at the cost of object separation. SAM3 demonstrates improved robustness across imaging conditions, particularly with noisy images. Third, ultralow-dose imaging enables substantial gains in acquisition speed without fundamentally limiting segmentation performance, provided that appropriate models were selected. These findings establish that effective model selection requires balancing segmentation accuracy, computational efficiency, robustness, and measurement fidelity. The framework presented here enables this selection process in a reproducible and application driven manner. While demonstrated for bacterial cell envelope thickness analysis, this approach is broadly applicable to microscopy workflows in which segmentation serves an intermediate step for quantitative measurement. Future work will focus on integrating this framework into automated acquisition pipelines to enable real-time, task-aware model selection during imaging.

## Availability of Data and Materials

The data underlying this article are available in Constellation at https://dx.doi.org/10.13139/ORNLNCCS/3025228. The models and framework scripts underlying this article are available on GitHub at https://github.com/Lynnicia/CryoEM_membranes_top_model_decision_tree and https://github.com/Sireesiru/Semantic-Segmentation-of-bacterial-cell-envelope-using-U-Nets. Fine-tuned SAM3 (Meta AI) models underlying this article are available on Hugging Face at https://huggingface.co/LynnMass/1024-SAM3-LD, https://huggingface.co/LynnMass/640-SAM3-LD, https://huggingface.co/LynnMass/1024-SAM3-ULD and https://huggingface.co/LynnMass/640-SAM-ULD.

## Acknowledgements

This work is supported by the U.S. Department of Energy, Office of Science FWP ERKCZ64, Structure Guided Design of Materials to Optimize the Abiotic-Biotic Material Interface, as part of the Biopreparedness Research Virtual Environment (BRaVE) initiative. Sample preparation, imaging and image analysis were conducted as part of a user project at the Center for Nanophase Materials Sciences (CNMS), which is a US Department of Energy, Office of Science User Facility at Oak Ridge National Laboratory. Electron microscopy data was collected using instrumentation within ORNL’s Materials Characterization Core provided by UT-Battelle, LLC, under Contract No. DE-AC05-00OR22725 with the DOE and sponsored by the Laboratory Directed Research and Development Program of Oak Ridge National Laboratory, managed by UT-Battelle, LLC, for the U.S. Department of Energy.

## Author Contributions Statement

L.N.M. and A.N.W. collected the cryoEM images, as well as designed, wrote and edited the manuscript. L.N.M. and S.S.M annotated the images, curated the models and evaluated model performance. S.R.B, A.N.B, C.R.H and J.L.M-F. prepared bacteria samples and BCET tool analysis. L.X.Z. and K.P. assisted with imaging and annotation with the BCET tool. R.K.V. and S.T.R. reviewed and edited the manuscript.

## Financial Support

This work is supported by the U.S. Department of Energy, Office of Science FWP ERKCZ64, Structure Guided Design of Materials to Optimize the Abiotic-Biotic Material Interface, as part of the Biopreparedness Research Virtual Environment (BRaVE) initiative. Sample preparation, imaging and image analysis were conducted as part of a user project at the Center for Nanophase Materials Sciences (CNMS), which is a US Department of Energy, Office of Science User Facility at Oak Ridge National Laboratory.

## Conflict of interest

The author(s) declare none

## Notes

https://github.com/Lynnicia/CryoEM_membranes_top_model_decision_tree

https://github.com/Sireesiru/Semantic-Segmentation-of-bacterial-cell-envelope-using-U-Nets

https://huggingface.co/LynnMass/1024-SAM3-LD

https://huggingface.co/LynnMass/640-SAM3-LD

https://huggingface.co/LynnMass/1024-SAM3-ULD

https://huggingface.co/LynnMass/640-SAM-ULD

https://dx.doi.org/10.13139/ORNLNCCS/3025228

### Competing Interest Statement

The authors have declared no competing interest.

## References

Archit, A., Freckmann, L., Nair, S., Khalid, N., Hilt, P., Rajashekar, V., Freitag, M., Teuber, C., Spitzner, M., Tapia Contreras, C., Buckley, G., von Haaren, S., Gupta, S., Grade, M., Wirth, M., Schneider, G., Dengel, A., Ahmed, S., & Pape, C. (2025). Segment Anything for Microscopy. Nature Methods, 22(3), 579–591. 10.1038/s41592-024-02580-4

Baumgartner, M., Jäger, P. F., Isensee, F., & Maier-Hein, K. H. (2021, 2021//). nnDetection: A Self-configuring Method for Medical Object Detection. Medical Image Computing and Computer Assisted Intervention – MICCAI 2021, Cham.

Bepler, T., Kelley, K., Noble, A. J., & Berger, B. (2020). Topaz-Denoise: general deep denoising models for cryoEM and cryoET. Nature Communications, 11(1), 5208. 10.1038/s41467-020-18952-1

Bible, A. N., Fletcher, S. J., Pelletier, D. A., Schadt, C. W., Jawdy, S. S., Weston, D. J., Engle, N. L., Tschaplinski, T., Masyuko, R., Polisetti, S., Bohn, P. W., Coutinho, T. A., Doktycz, M. J., & Morrell-Falvey, J. L. (2016). A Carotenoid-Deficient Mutant in Pantoea sp. YR343, a Bacteria Isolated from the Rhizosphere of Populus deltoides, Is Defective in Root Colonization [Original Research]. Frontiers in Microbiology, *Volume* 7 - 2016. 10.3389/fmicb.2016.00491

Bishop, C. (2007). *Pattern Recognition and Machine Learning (Information Science and Statistics)*.

Buchholz, T. O., Jordan, M., Pigino, G., & Jug, F. (2019, 8-11 April 2019). Cryo-CARE: Content-Aware Image Restoration for Cryo-Transmission Electron Microscopy Data. 2019 IEEE 16th International Symposium on Biomedical Imaging (ISBI 2019),

Carion, N., Gustafson, L., Hu, Y.-T., Debnath, S., Hu, R., Suris, D., Ryali, C., Alwala, K. V., Khedr, H., & Huang, A. (2025). Sam 3: Segment anything with concepts. *arXiv preprint arXiv*:2511.16719.

Chen, L., Shen, J., Li, X., Li, R., Gao, X., Chen, X., Pan, X., & Jin, X. (2025). Utilizing Detectron2 for accurate and efficient colon cancer detection in histopathological images. Front Bioeng Biotechnol, 13, 1593534. 10.3389/fbioe.2025.1593534

Chen, T., Cao, R., Yu, X., Zhu, L., Ding, C., Ji, D., Chen, C., Zhu, Q., Xu, C., & Mao, P. (2025). SAM3-Adapter: Efficient Adaptation of Segment Anything 3 for Camouflage Object Segmentation, Shadow Detection, and Medical Image Segmentation. *arXiv preprint arXiv*:2511.19425.

Ciaglia, F., Zuppichini, F. S., Guerrie, P., McQuade, M., & Solawetz, J. (2022). Roboflow 100: A rich, multi-domain object detection benchmark. *arXiv preprint arXiv*:2211.13523.

Egerton, R. (2020). Understanding Radiation Damage in Beam-Sensitive TEM Specimens. Microscopy and Microanalysis, 26(S2), 84–86. 10.1017/s1431927620013331

Everingham, M., Van Gool, L., Williams, C. K. I., Winn, J., & Zisserman, A. (2010). The Pascal Visual Object Classes (VOC) Challenge. International Journal of Computer Vision, 88(2), 303–338. 10.1007/s11263-009-0275-4

Halsted, M. C., Bible, A. N., Morrell-Falvey, J. L., & Retterer, S. T. (2022). Quantifying biofilm propagation on chemically modified surfaces. Biofilm, 4, 100088. 10.1016/j.bioflm.2022.100088

He, K., Gkioxari, G., Dollár, P., & Girshick, R. (2017). Mask r-cnn. Proceedings of the IEEE international conference on computer vision,

He, S., Bao, R., Li, J., Stout, J., Bjornerud, A., Grant, P. E., & Ou, Y. (2023). Computer-vision benchmark segment-anything model (sam) in medical images: Accuracy in 12 datasets. *arXiv preprint arXiv*:2304.09324.

Israel, U., Marks, M., Dilip, R., Li, Q., Yu, C., Laubscher, E., Iqbal, A., Pradhan, E., Ates, A., Abt, M., Brown, C., Pao, E., Li, S., Pearson-Goulart, A., Perona, P., Gkioxari, G., Barnowski, R., Yue, Y., & Van Valen, D. (2025). CellSAM: A Foundation Model for Cell Segmentation. bioRxiv. 10.1101/2023.11.17.567630

Jiang, C., Ding, T., Song, C., Tu, J., Yan, Z., Shao, Y., Wang, Z., Shang, Y., Han, T., & Tian, Y. (2026). Medical SAM3: A Foundation Model for Universal Prompt-Driven Medical Image Segmentation. *arXiv preprint arXiv*:2601.10880.

Jocher, G., & Qiu, J. (2024). Ultralytics yolo11. 2024. In.

Kazimi, B., & Sandfeld, S. (2024). Enhancing Semantic Segmentation in High-Resolution TEM Images: A Comparative Study of Batch Normalization and Instance Normalization. Microscopy and Microanalysis, 31(1). 10.1093/mam/ozae093

Khanam, R., & Hussain, M. (2024). Yolov11: An overview of the key architectural enhancements. *arXiv preprint arXiv*:2410.17725.

Kirillov, A., Mintun, E., Ravi, N., Mao, H., Rolland, C., Gustafson, L., Xiao, T., Whitehead, S., Berg, A. C., & Lo, W.-Y. (2023). Segment anything. Proceedings of the IEEE/CVF international conference on computer vision,

Kuijper, M., van Hoften, G., Janssen, B., Geurink, R., De Carlo, S., Vos, M., van Duinen, G., van Haeringen, B., & Storms, M. (2015). FEI’s direct electron detector developments: Embarking on a revolution in cryo-TEM. J Struct Biol, 192(2), 179–187. 10.1016/j.jsb.2015.09.014

Lin, T.-Y., Maire, M., Belongie, S., Hays, J., Perona, P., Ramanan, D., Dollár, P., & Zitnick, C. L. (2014). Microsoft coco: Common objects in context. European conference on computer vision,

Long, J., Shelhamer, E., & Darrell, T. (2015, 7-12 June 2015). Fully convolutional networks for semantic segmentation. 2015 IEEE Conference on Computer Vision and Pattern Recognition (CVPR),

Madugula, S. S., Massenburg, L. N., Brown, S. R., Bible, A. N., Harris, C. R., Zhang, L. X., Parker, K., Retterer, S. T., Morrell-Falvey, J. L., Vasudevan, R. K., & Williams, A. N. (2026). Automated Bacterial Identification and Morphological Feature Analysis in Low-Dose Cryo-EM Using YOLOv11. Advanced Intelligent Discovery, *n/a*(n/a), e202500241. 10.1002/aidi.202500241

Manning, C. D., Raghavan, P., & Schütze, H. (2008). Introduction to information retrieval.

Mendez, J. H., Mehrani, A., Randolph, P., & Stagg, S. (2019). Throughput and resolution with a next-generation direct electron detector. IUCrJ, 6(Pt 6), 1007–1013. 10.1107/s2052252519012661

Odena, A., Dumoulin, V., & Olah, C. (2016). Deconvolution and checkerboard artifacts. Distill, 1(10), e3.

Ronneberger, O., Fischer, P., & Brox, T. (2015). U-Net: Convolutional Networks for Biomedical Image Segmentation. In N. Navab, J. Hornegger, W. M. Wells, & A. F. Frangi, Medical Image Computing and Computer-Assisted Intervention – MICCAI 2015 Cham.

Sanchez-Garcia, R., Gomez-Blanco, J., Cuervo, A., Carazo, J. M., Sorzano, C. O. S., & Vargas, J. (2021). DeepEMhancer: a deep learning solution for cryo-EM volume post-processing. Communications Biology, 4(1), 874. 10.1038/s42003-021-02399-1

Sapkota, R., Cheppally, R. H., Sharda, A., & Karkee, M. (2025). YOLO26: key architectural enhancements and performance benchmarking for real-time object detection. *arXiv preprint arXiv*:2509.25164.

Sapkota, R., & Karkee, M. (2025). Ultralytics YOLO evolution: An overview of YOLO26, YOLO11, YOLOv8 and YOLOv5 object detectors for computer vision and pattern recognition. *arXiv preprint arXiv*:2510.09653.

Shelhamer, E., Long, J., & Darrell, T. (2017). Fully Convolutional Networks for Semantic Segmentation. IEEE Transactions on Pattern Analysis and Machine Intelligence, 39(4), 640–651. 10.1109/TPAMI.2016.2572683

Stringer, C., Wang, T., Michaelos, M., & Pachitariu, M. (2021). Cellpose: a generalist algorithm for cellular segmentation. Nature Methods, 18(1), 100–106. 10.1038/s41592-020-01018-x

Sudre, C. H., Li, W., Vercauteren, T., Ourselin, S., & Jorge Cardoso, M. (2017). Generalised Dice Overlap as a Deep Learning Loss Function for Highly Unbalanced Segmentations. In M. J. Cardoso, T. Arbel, G. Carneiro, T. Syeda-Mahmood, J. M. R. S. Tavares, M. Moradi, A. Bradley, H. Greenspan, J. P. Papa, A. Madabhushi, J. C. Nascimento, J. S. Cardoso, V. Belagiannis, & Z. Lu, Deep Learning in Medical Image Analysis and Multimodal Learning for Clinical Decision Support Cham.

Sugawara, Y., Shiota, S., & Kiya, H. (2019). Checkerboard artifacts free convolutional neural networks. APSIPA Transactions on Signal and Information Processing, 8, e9, Article e9. 10.1017/ATSIP.2019.2

Taghanaki, S. A., Zheng, Y., Kevin Zhou, S., Georgescu, B., Sharma, P., Xu, D., Comaniciu, D., & Hamarneh, G. (2019). Combo loss: Handling input and output imbalance in multi-organ segmentation. Computerized Medical Imaging and Graphics, 75, 24–33. 10.1016/j.compmedimag.2019.04.005

Travieso, G., Benatti, A., & Costa, L. D. F. (2024). An analytical approach to the Jaccard similarity index. *arXiv preprint arXiv*:2410.16436.

Wang, A., Xu, M., Goel, R., Shabeeb, Z., Panicker, I., & Jamali, V. (2025). SAM-EM: Real-Time Segmentation for Automated Liquid Phase Transmission Electron Microscopy. *arXiv preprint arXiv*:2501.03153.

Wang, X., Howe, S., Deng, F., & Zhao, J. (2021). Current Applications of Absolute Bacterial Quantification in Microbiome Studies and Decision-Making Regarding Different Biological Questions. Microorganisms, 9(9). 10.3390/microorganisms9091797

Zou, K. H., Warfield, S. K., Bharatha, A., Tempany, C. M., Kaus, M. R., Haker, S. J., Wells, W. M., 3rd, Jolesz, F. A., & Kikinis, R. (2004). Statistical validation of image segmentation quality based on a spatial overlap index. Acad Radiol, 11(2), 178–189. 10.1016/s1076-6332(03)00671-8

