## Supplementary Information for "Top Model Decision Tree: Selecting Segmentation Models for Reliable Quantitative Analysis in Low- and Ultralow-Dose CryoEM"

**Supplementary Figures**

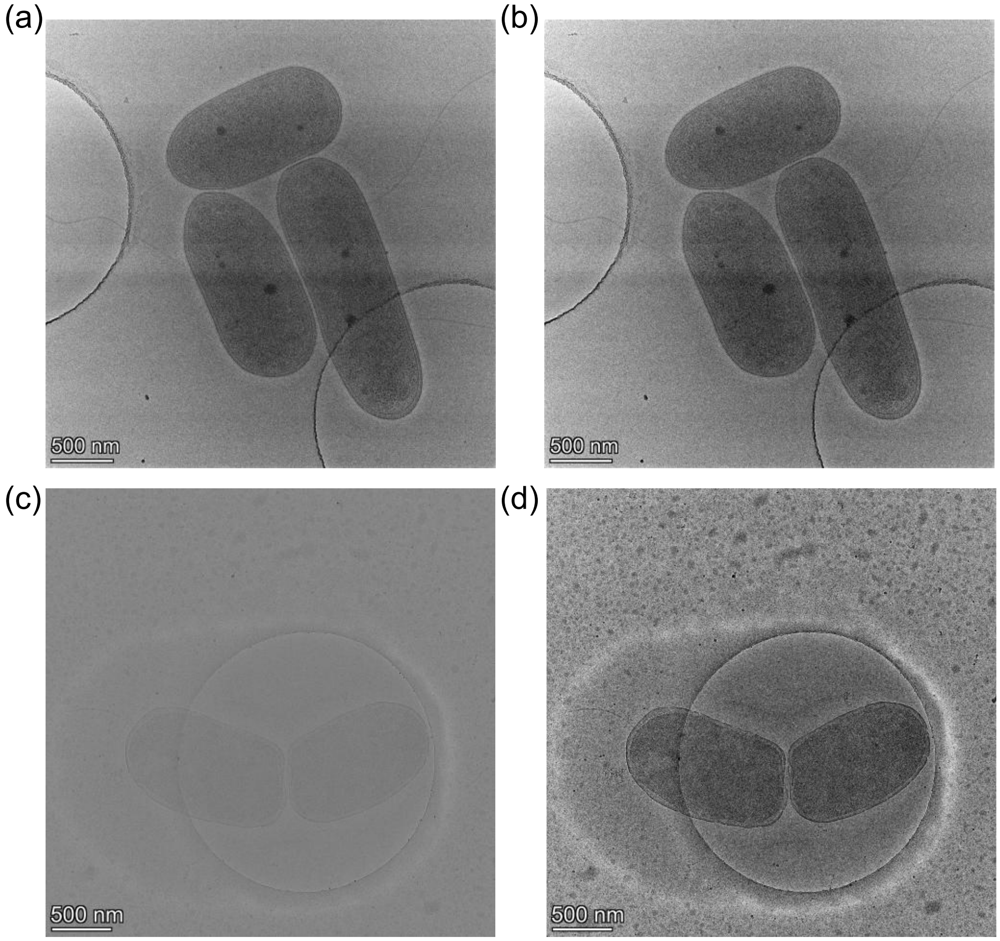


**Supplementary Fig. 1.** Example dynamic normalization on low-dose and ultralow-dose images. (a) Raw and (b) dynamically normalized low-dose cryoEM images. (c) Raw and (d) dynamically normalized ultralow-dose cryoEM images. Scale bar is 500 nm.

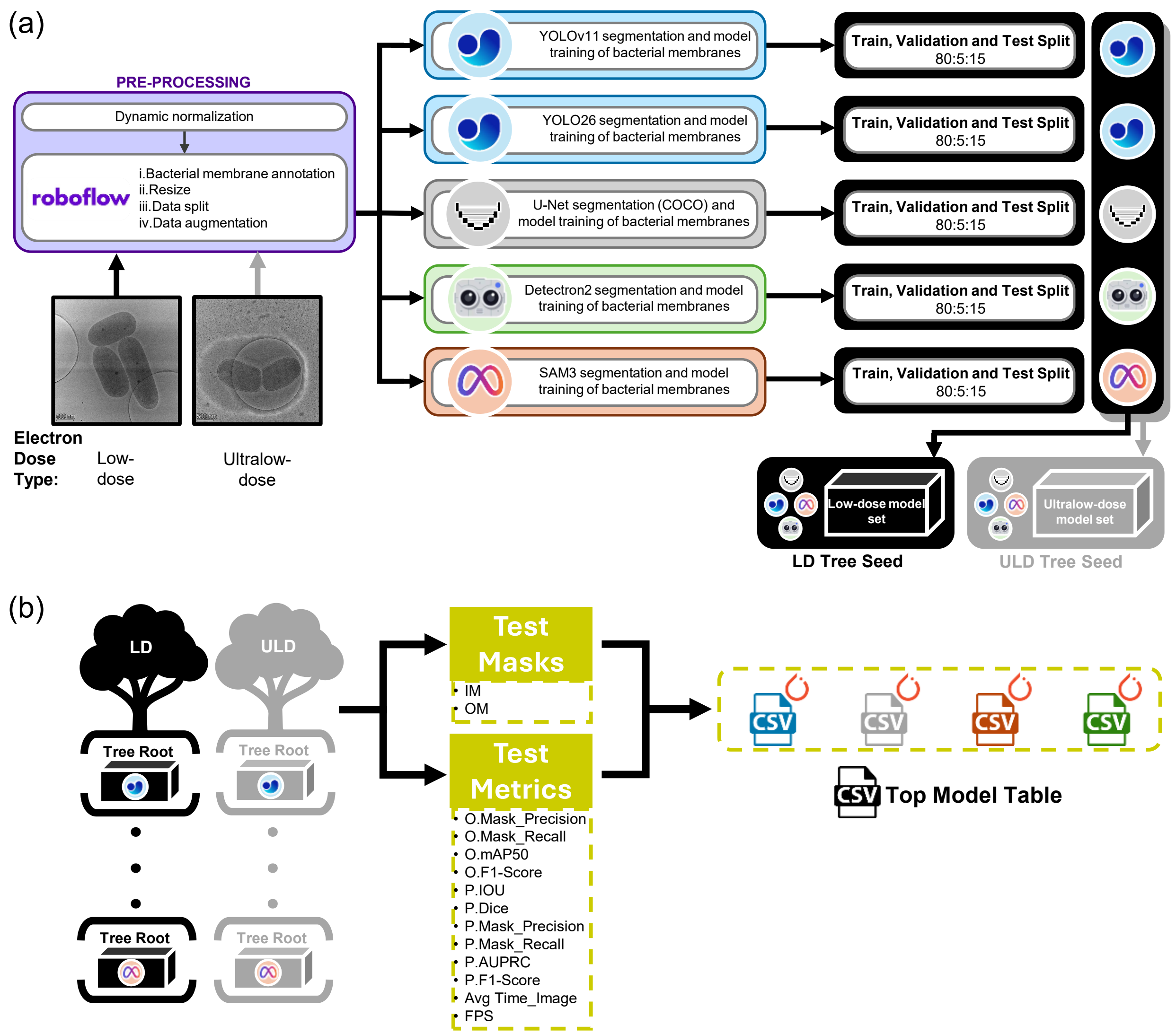


**Supplementary Fig. 2.** Workflow to generate and utilize low-dose and ultra-low dose model seeds. (a) Low-dose (LD) and ultralow-dose (ULD) images were separately pre-processed and used to fine-tune YOLOv11, YOLO26, U-Net, Detectron2 and SAM3 base models at an 80:5:15 train-valid-test split to generate the model seeds. The test images and annotations were used for unbiased test metrics and masks. Test images can be generated independently from the split workflow. (b) Utilization of model seeds was designed to output test IM and OM masks and test metrics for unbiased test images and annotations. "O." signifies overall best metrics based on the 0.5 IoU threshold for instance segmentations and overall best metrics for semantic segmentations across the confidence thresholds. "P." signifies per-image mean macro metrics. Either the LD seed, the ULD seed or both seeds can be selected for test image masks and metrics.

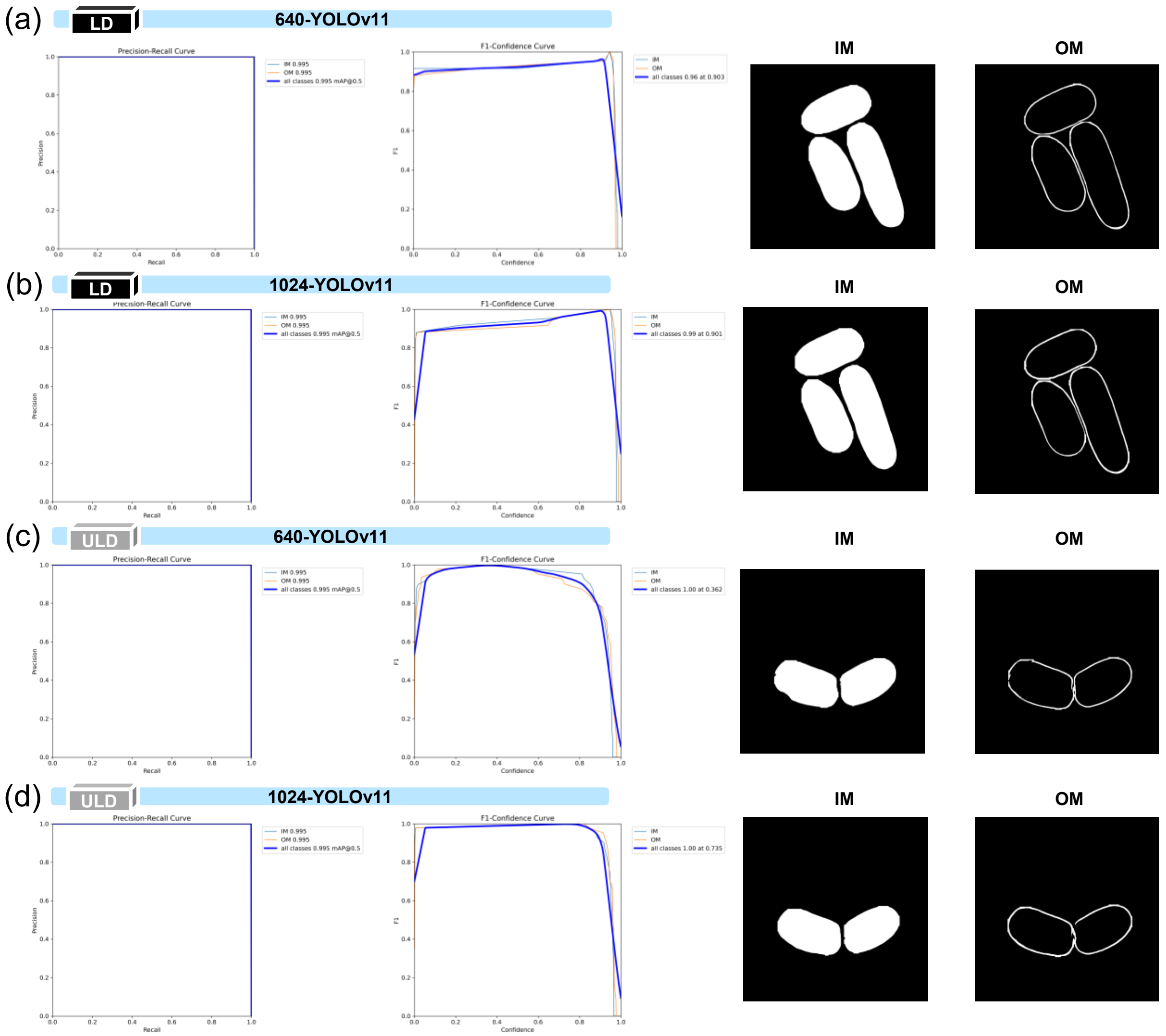


**Supplementary Fig. 3.** Valid image evaluation metrics curves and test image-extracted masks for four YOLOv11 models. From left to right, PR curves, F1 curves, inner membrane (IM) binary mask and outer membrane (OM) binary mask for (a) 640-YOLOv11 and (b) 1024-YOLOv11 models trained on the LD seed (low-dose images), as well as (c) 640-YOLOv11 and (d) 1024-YOLOv11 models trained on the ULD seed (ultralow-dose images). The values 640 and 1024 refer to the train image size for fine-tuning the pre-trained base YOLOv11 instance segmentation model. The 640-YOLOv11-LD model is sourced from Madugula et al. (Madugula et al., 2026).

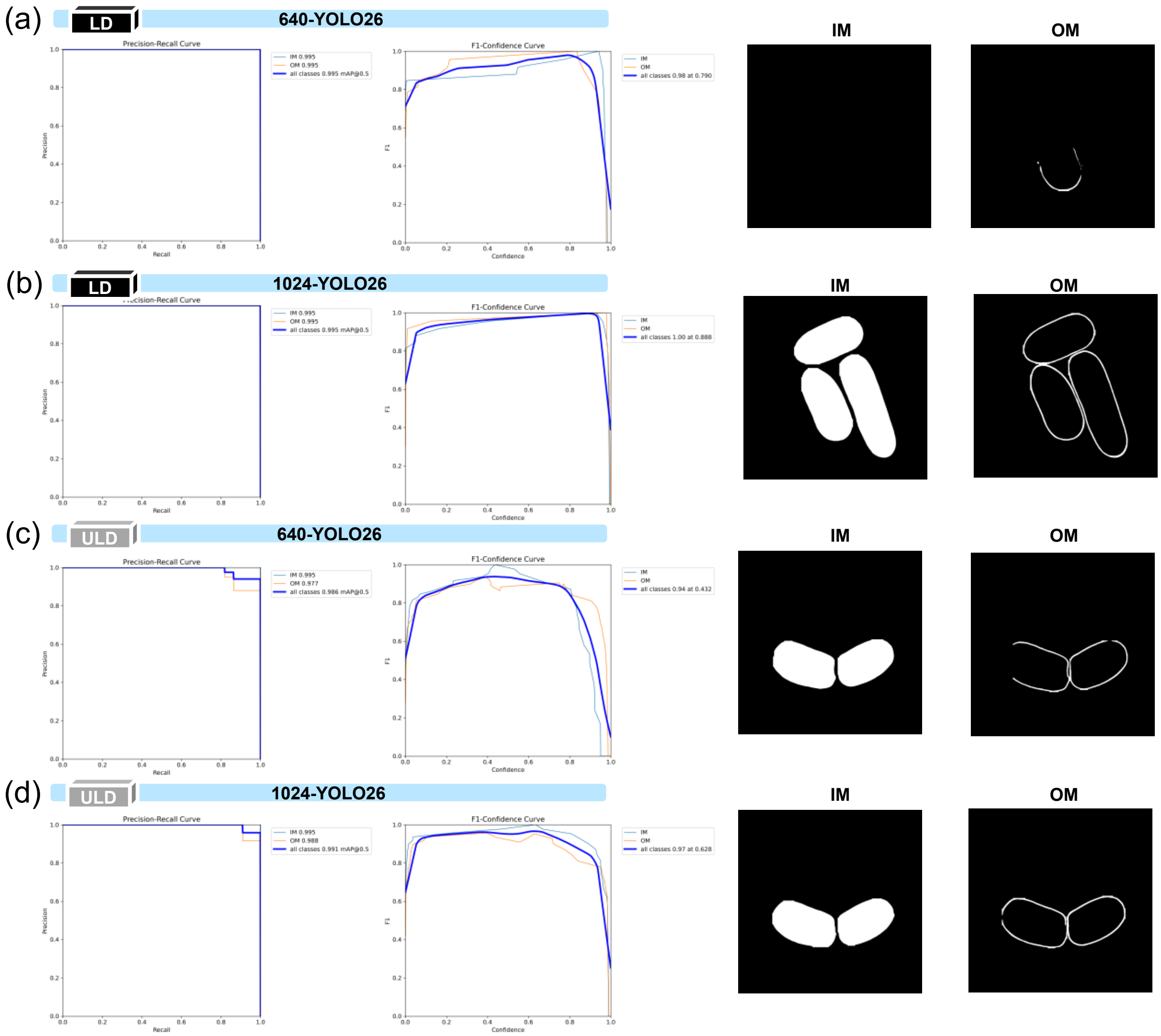


**Supplementary Fig. 4.** Valid image evaluation metrics curves and test image-extracted masks for four YOLO26 models. From left to right, PR curves, F1 curves, inner membrane (IM) binary mask and outer membrane (OM) binary mask for (a) 640-YOLO26 and (b) 1024-YOLO26 models trained on the LD seed (low-dose images), as well as (c) 640-YOLO26 and (d) 1024-YOLO26 models trained on the ULD seed (ultralow-dose images). The values 640 and 1024 refer to the train image size for fine-tuning the pre-trained base YOLO26 instance segmentation model.

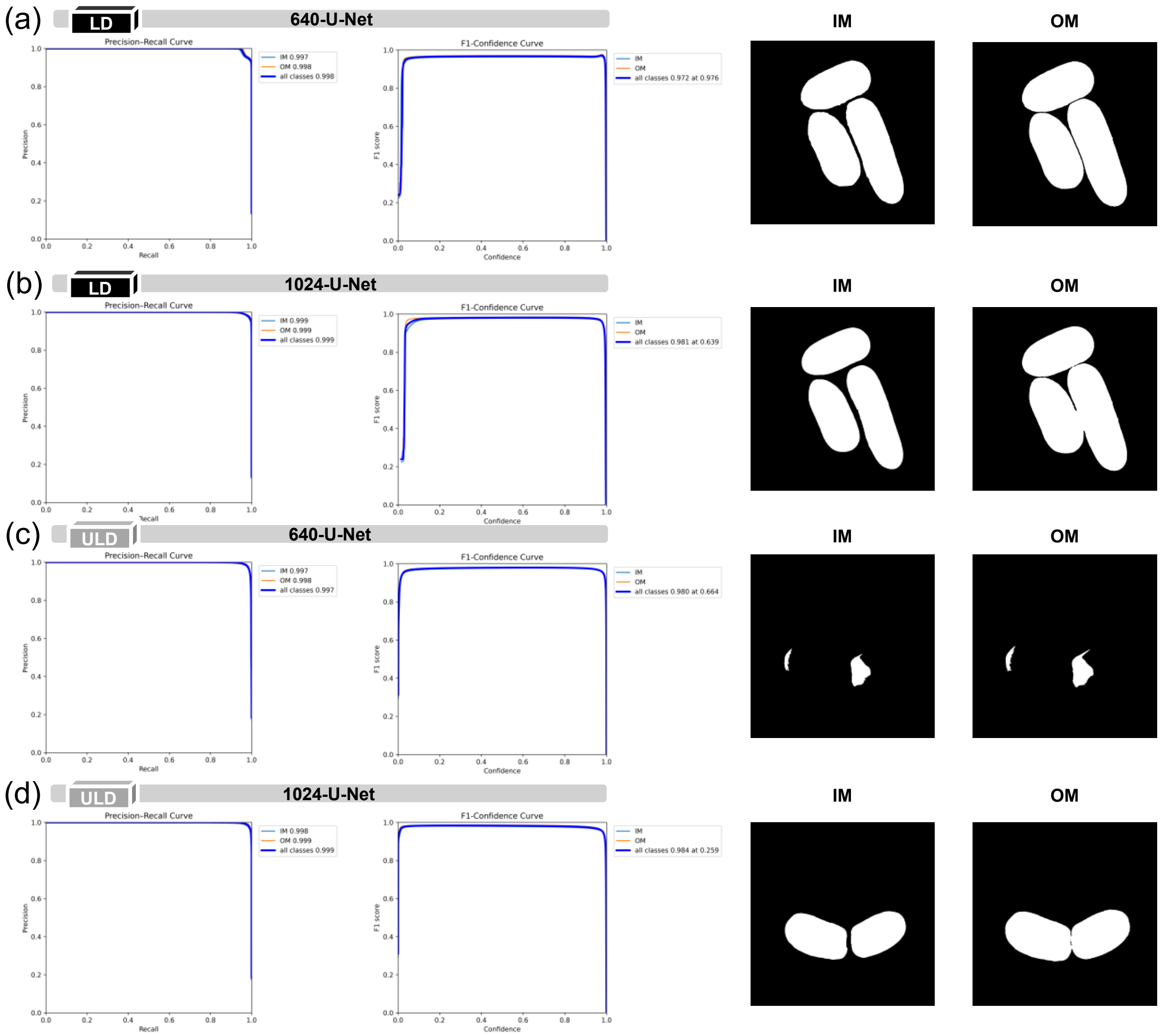


**Supplementary Fig. 5.** Valid image evaluation metrics curves and test image-extracted masks for four U-Net models. From left to right, PR curves, F1 curves, inner membrane (IM) binary mask and outer membrane (OM) binary mask for (a) 640-U-Net and (b) 1024-U-Net models trained on the LD seed (low-dose images), as well as (c) 640-U-Net and (d) 1024-U-Net models trained on the ULD seed (ultralow-dose images). The values 640 and 1024 refer to the train image size for fine-tuning the pre-trained base U-Net semantic segmentation model.

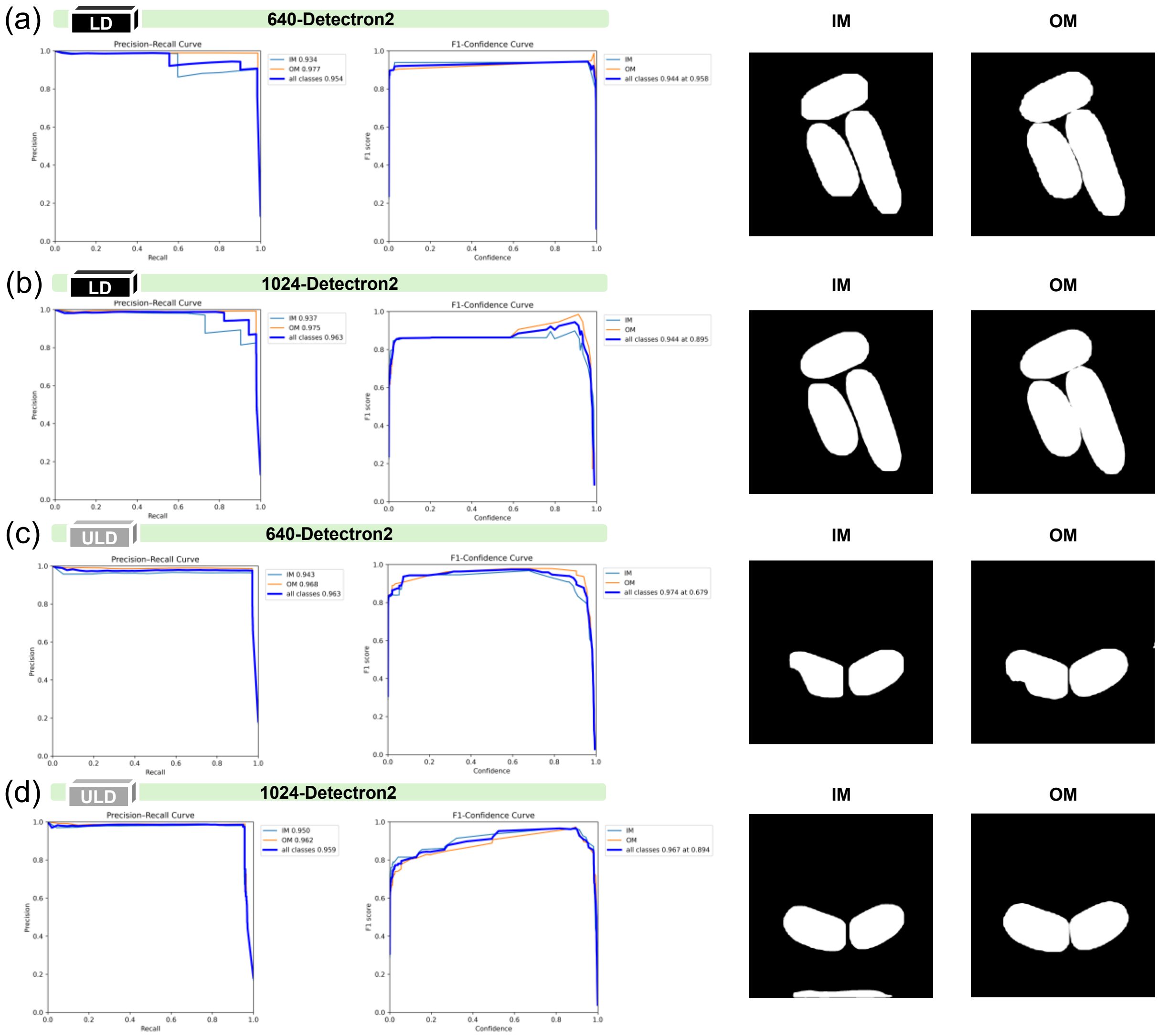


**Supplementary Fig. 6.** Valid image evaluation metrics curves and test image-extracted masks for four Detectron2 models. From left to right, PR curves, F1 curves, inner membrane (IM) binary mask and outer membrane (OM) binary mask for (a) 640-Detectron2 and (b) 1024-Detectron2 models trained on the LD seed (low-dose images), as well as (c) 640-Detectron2 and (d) 1024-Detectron2 models trained on the ULD seed (ultralow-dose images). The values 640 and 1024 refer to the train image size for fine-tuning the pre-trained base Detectron2 instance segmentation model.

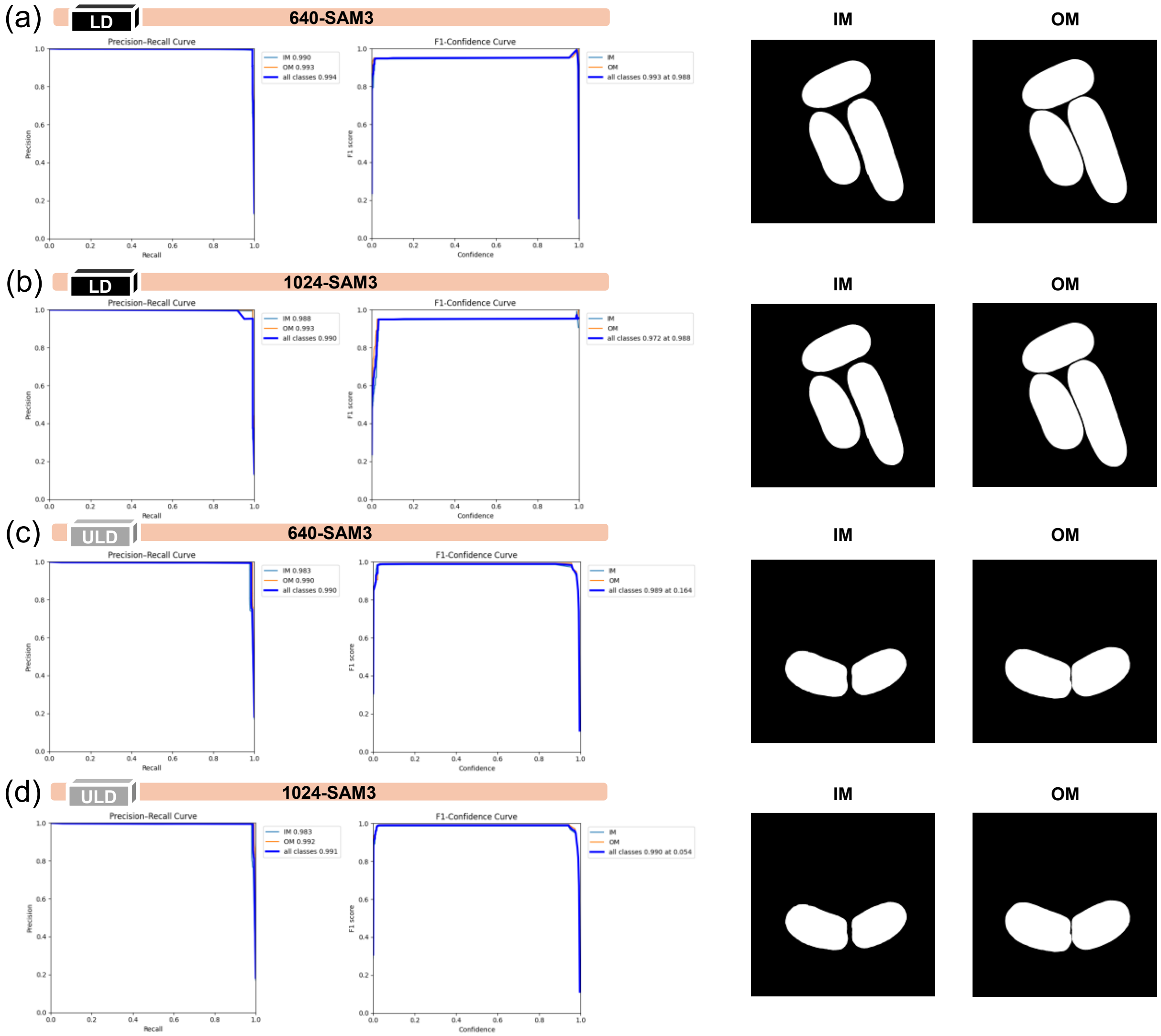


**Supplementary Fig. 7.** Valid image evaluation metrics curves and test image-extracted masks for four SAM3 models. From left to right, PR curves, F1 curves, inner membrane (IM) binary mask and outer membrane (OM) binary mask for (a) 640-SAM3 and (b) 1024-SAM3 models trained on the LD seed (low-dose images), as well as (c) 640-SAM3 and (d) 1024-SAM3 models trained on the ULD seed (ultralow-dose images). The values 640 and 1024 refer to the train image size for fine-tuning the pre-trained base SAM3 instance segmentation model.

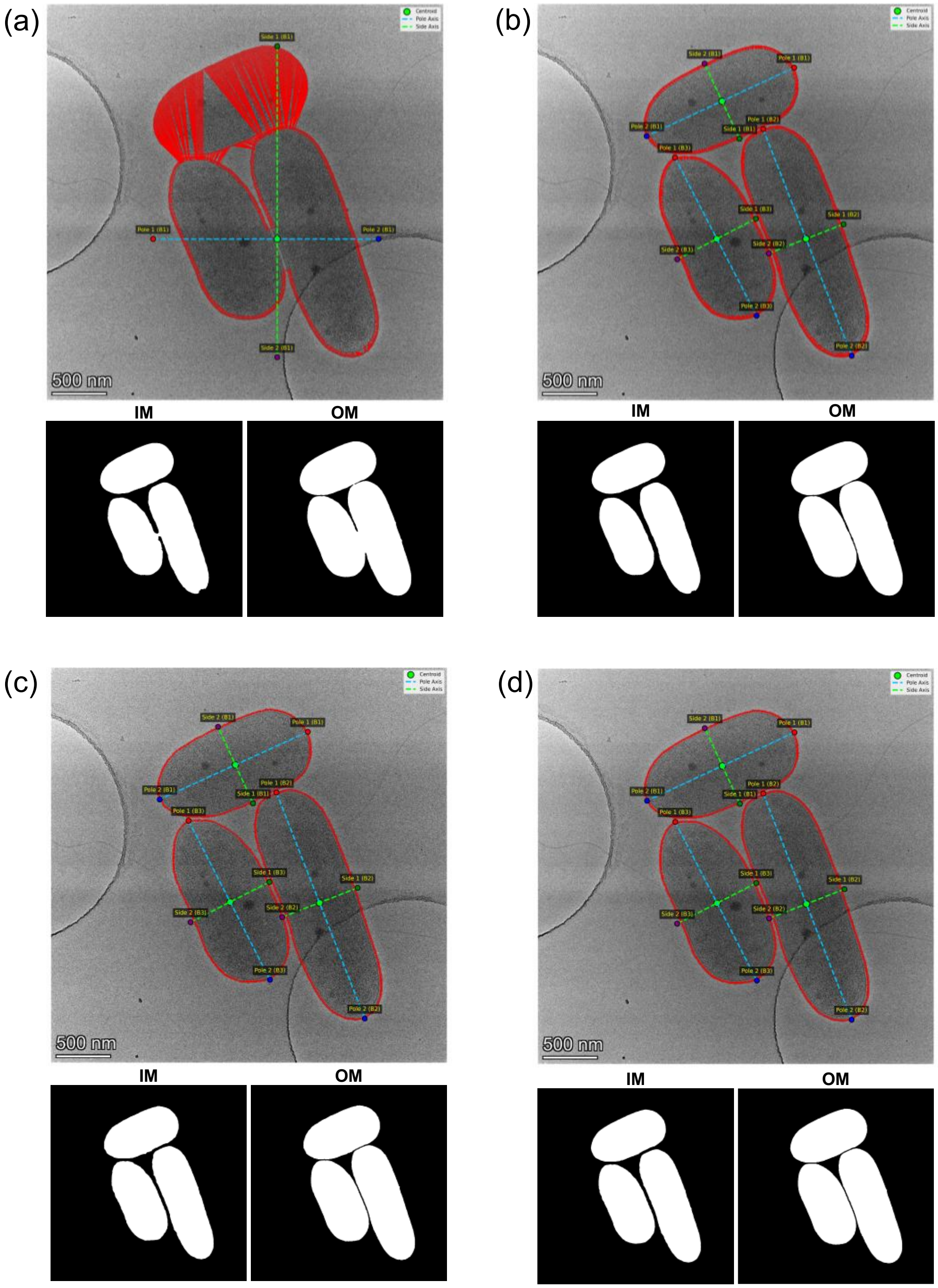


**Supplementary Fig. 8.** Image interpolation effects on extracted binary masks. (a) Inter-linear interpolation and (b) inter-area interpolation to downsize low-dose test images from 4096 × 4096 to 1024 × 1024 for mask generation on the 1024-U-Net-LD model. (c) Inter-linear interpolation and (d) inter-area interpolation to downsize low-dose test images from 4096 × 4096 to 1024 × 1024 for mask generation on the 640-U-Net-LD model.

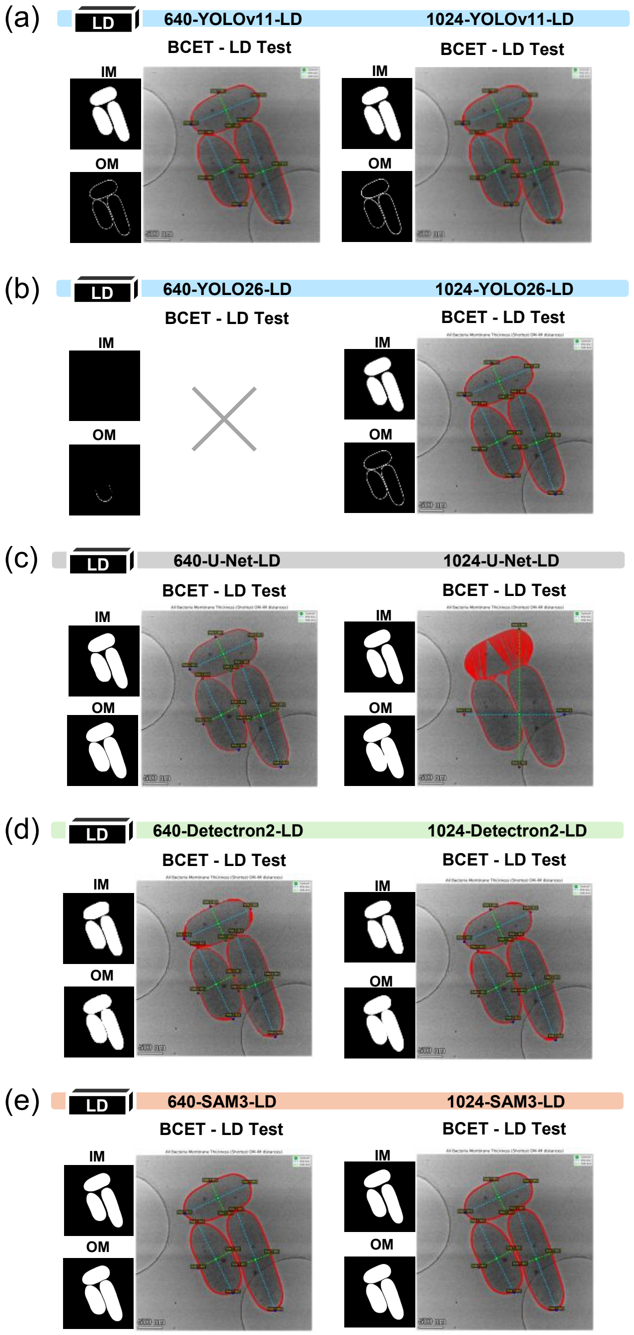


**Supplementary Fig. 9.** Example BCET outputs and corresponding binary IM and OM masks for models evaluated within the LD model seed. BCET envelope thickness measurements are shown for (a) 640-YOLOv11-LD and 1024-YOLOv11-LD, (b) 640-YOLO26-LD and 1024-YOLO26-LD, (c) 640-U-Net-LD and 1024-U-Net-LD, (d) 640-Detectron2-LD and 1024-Detectron2-LD, and (e) 640-SAM3-LD and 1024-SAM3-LD. Models were evaluated on low-dose test images to illustrate differences in mask behavior and BCET measurement outcomes. Scale bar is 500 nm.

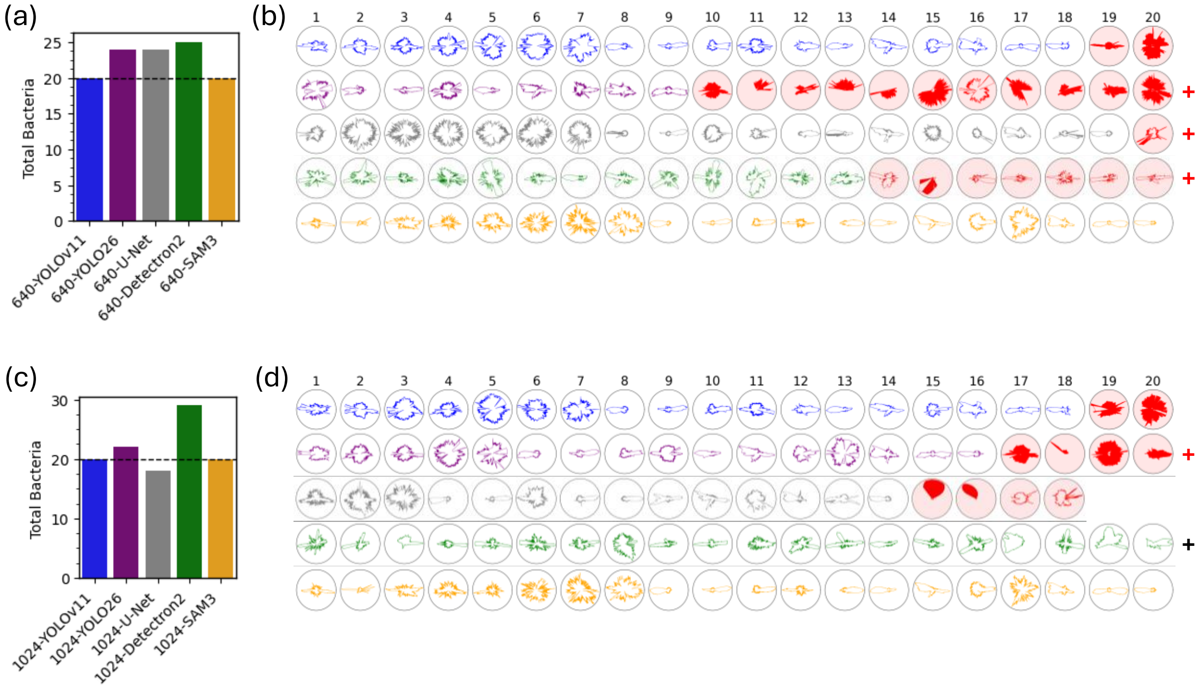


**Supplementary Fig. 10.** Mask screening reveals biologically relevant model performance with low-dose test images. (a) Total bacteria count per model based on a ground truth of 20 bacteria. (b) Radial plots of the bacterial cell envelope thickness of each bacterium measured by the BCET tool with 640-LD models. A cutoff of 20 radial plots are depicted with models sorted by cell envelope thickness measurement outputs above 0 nm and no merged bacterial objects (left-to-right). Radial plots of bacteria with erroneous envelope thickness measurements of 0 nm were colored red (right). Plot colors represent YOLOv11 (blue), YOLO26 (purple), U-Net (grey), Detectron2 (green) and SAM3 (orange) models. (c) Total bacteria count and (d) radial plots of the BCET tool for 1024-LD models. A red plus sign indicates additional radial plots with 0 nm envelope thickness measurement outputs or merged bacterial neighbors, and a black plus sign indicates additional outputs for cell envelope thickness measurements at 0 nm (erroneous) and above 0 nm.

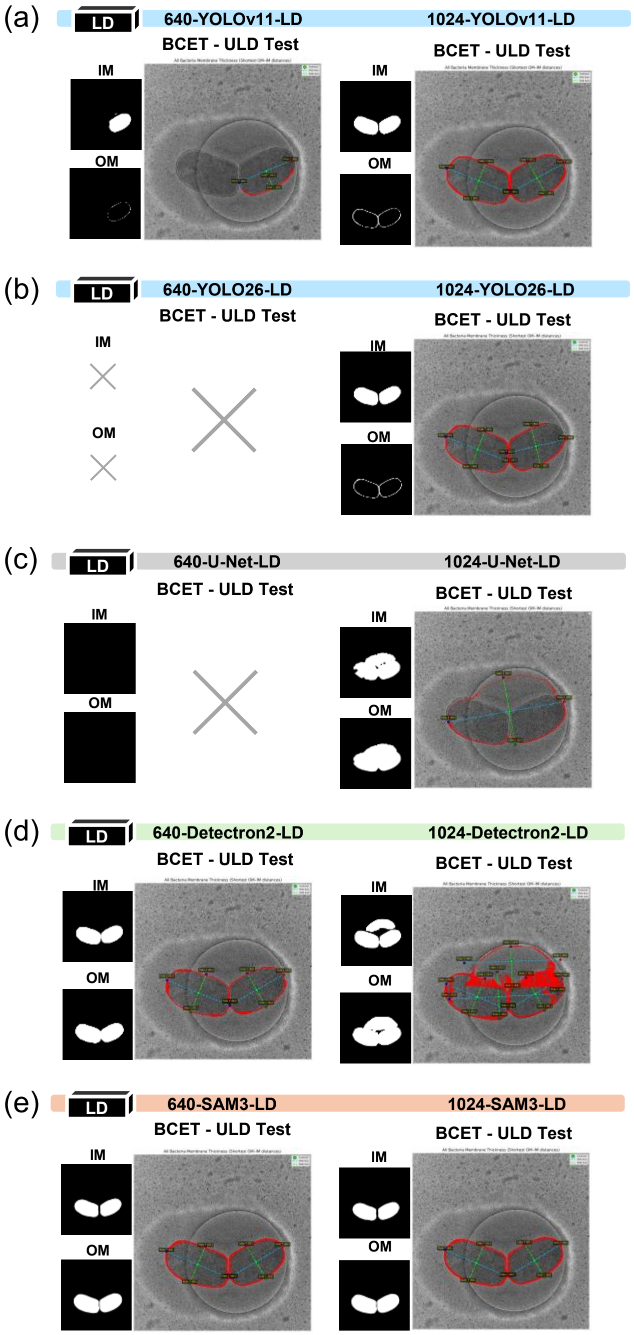


**Supplementary Fig. 11.** Example BCET outputs and corresponding binary IM and OM masks for models evaluated within the LD model seed. BCET envelope thickness measurements are shown for (a) 640-YOLOv11-LD and 1024-YOLOv11-LD, (b) 640-YOLO26-LD and 1024-YOLO26-LD, (c) 640-U-Net-LD and 1024-U-Net-LD, (d) 640-Detectron2-LD and 1024-Detectron2-LD, and (e) 640-SAM3-LD and 1024-SAM3-LD. Models were evaluated on ultralow-dose test images to illustrate differences in mask behavior and BCET measurement outcomes. Scale bar is 500 nm.

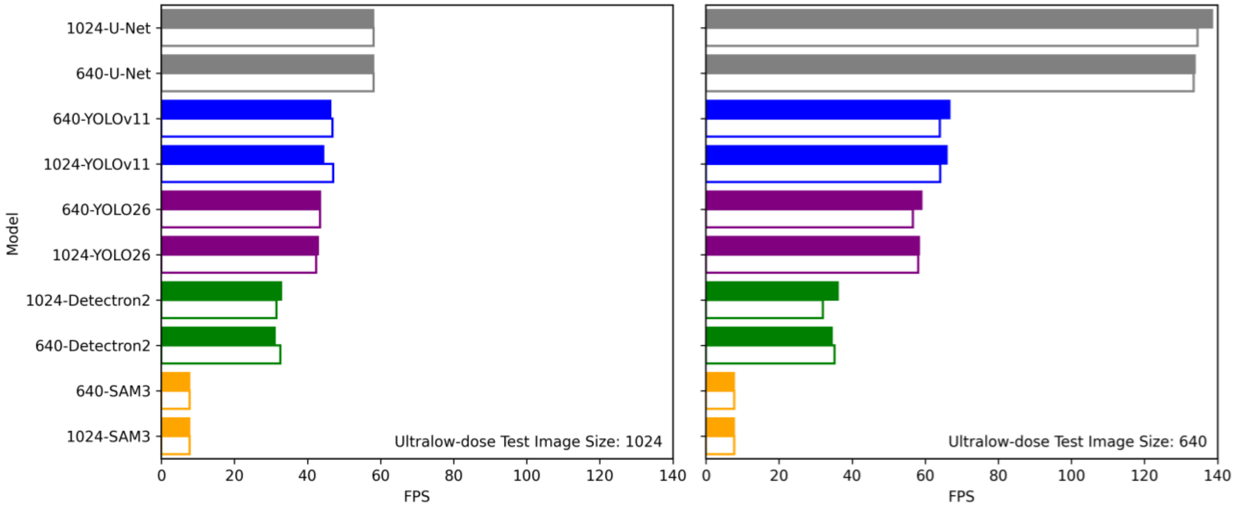


**Supplementary Fig. 12.** Bar graphs of FPS inferencing speed of the LD model seed and ULD model seed for ultralow-dose test images inferenced at 640 × 640. The models in the LD seed are represented as filled bars and models in the ULD seed are represented as open bars. Model seed train image sizes represented in the left plot (640 train image size) or the right plot (1024 train image size). Colors represent YOLOv11 (blue), YOLO26 (purple), U-Net (grey), Detectron2 (green) and SAM3 (orange).

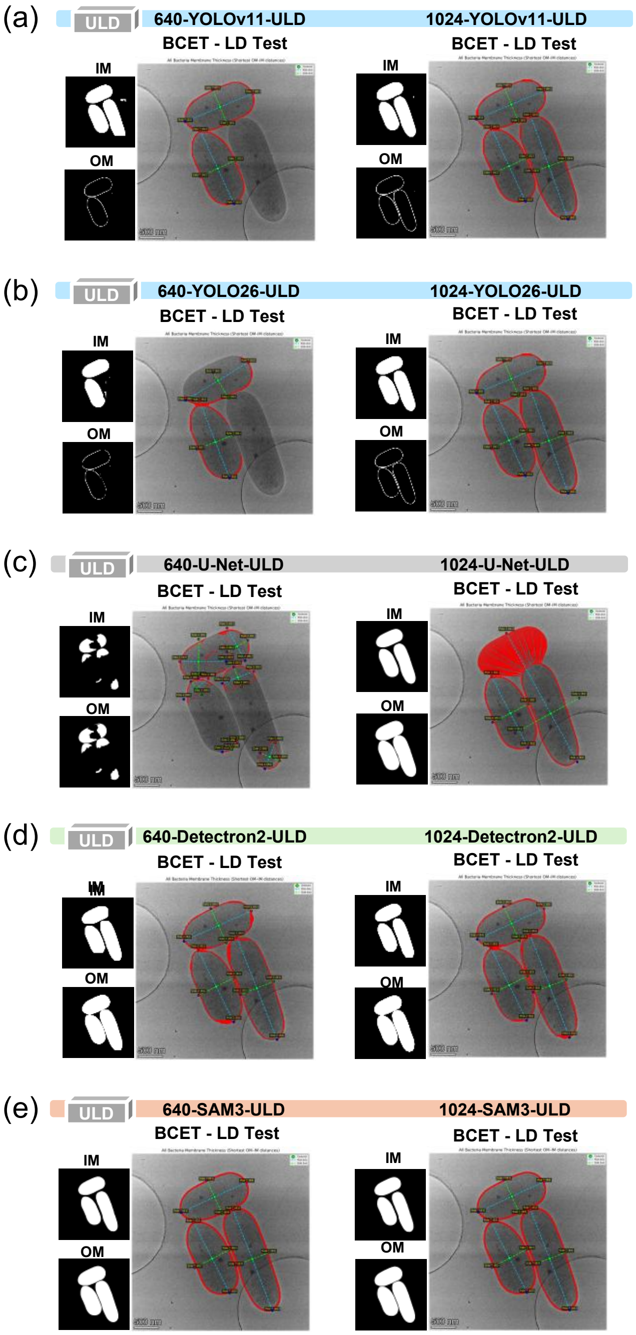


**Supplementary Fig. 13.** Example BCET outputs and corresponding binary IM and OM masks for models evaluated within the ULD model seed. BCET envelope thickness measurements are shown for (a) 640-YOLOv11-ULD and 1024-YOLOv11-ULD, (b) 640-YOLO26-ULD and 1024-YOLO26-ULD, (c) 640-U-Net-ULD and 1024-U-Net-ULD, (d) 640-Detectron2-ULD and 1024-Detectron2-ULD, and (e) 640-SAM3-ULD and 1024-SAM3-ULD. Models were evaluated on low-dose test images to illustrate differences in mask behavior and BCET measurement outcomes. Scale bar is 500 nm.

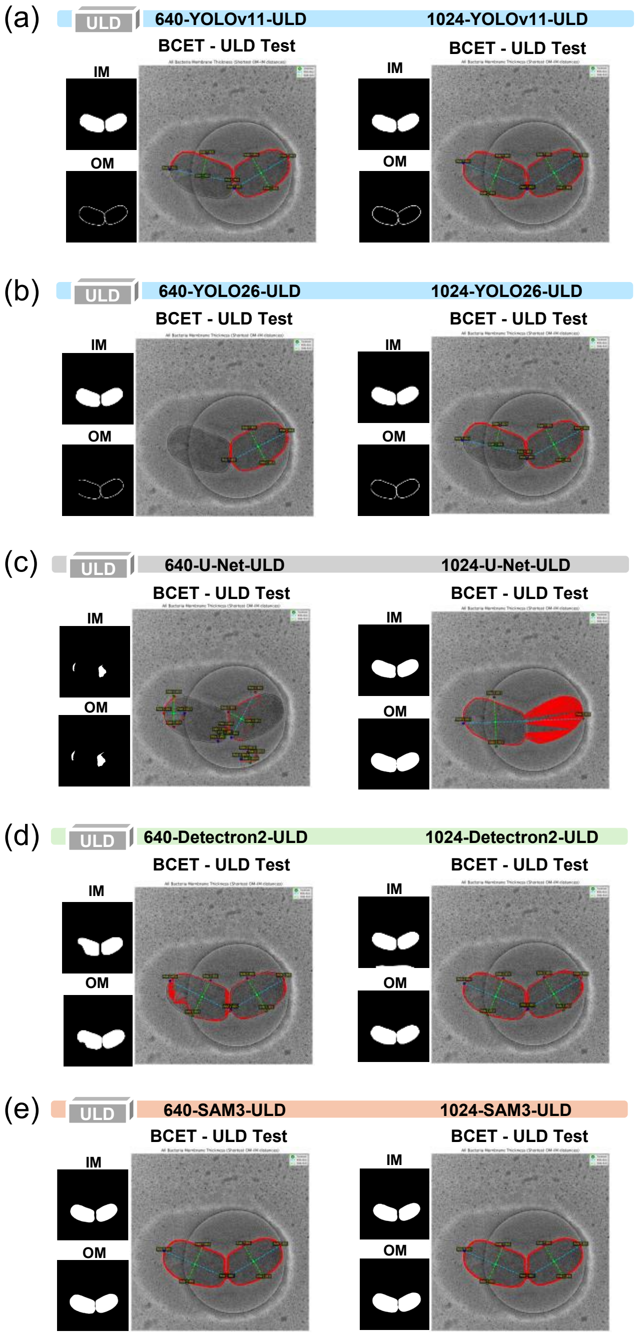


**Supplementary Fig. 14.** Example BCET outputs and corresponding binary IM and OM masks for models evaluated within the ULD model seed. BCET envelope thickness measurements are shown for (a) 640-YOLOv11-ULD and 1024-YOLOv11-ULD, (b) 640-YOLO26-ULD and 1024-YOLO26-ULD, (c) 640-U-Net-ULD and 1024-U-Net-ULD, (d) 640-Detectron2-ULD and 1024-Detectron2-ULD, and (e) 640-SAM3-ULD and 1024-SAM3-ULD. Models were evaluated on ultralow-dose test images to illustrate differences in mask behavior and BCET measurement outcomes. Scale bar is 500 nm.

### Supplementary Tables

**Supplementary Table 1.** Performance metrics on the valid images for the low-dose (LD) OM-IM fine-tuned models.

| Model | Electron Dose | Class | Mask Precision* | Mask Recall* | mAP50/ AUPRC* | Best F1-Score* | Mean IOU† | Mean Dice† | Mean Mask Precision† | Mean Mask Recall† | AUPRC† | Mean F1-Score† |
|---|---|---|---|---|---|---|---|---|---|---|---|---|
| **640-YOLOv11-LD** | LD | All | 0.914 | 1.0 | 0.995 | 0.955 | 0.834 | 0.903 | 0.889 | 0.922 | 0.898 | 0.903 |
| | | IM | 0.913 | 1.0 | 0.995 | 0.955 | 0.942 | 0.968 | 0.952 | 0.989 | 0.947 | 0.968 |
| | | OM | 0.915 | 1.0 | 0.995 | 0.956 | 0.726 | 0.838 | 0.826 | 0.856 | 0.820 | 0.838 |
| **1024-YOLOv11-LD** | | All | 0.985 | 1.0 | 0.995 | 0.992 | 0.843 | 0.909 | 0.910 | 0.915 | 0.889 | 0.909 |
| | | IM | 0.984 | 1.0 | 0.995 | 0.992 | 0.933 | 0.963 | 0.949 | 0.983 | 0.946 | 0.963 |
| | | OM | 0.985 | 1.0 | 0.995 | 0.993 | 0.753 | 0.855 | 0.871 | 0.846 | 0.832 | 0.855 |
| **640-YOLO26-LD** | | All | 0.961 | 1.0 | 0.995 | 0.980 | 0.801 | 0.880 | 0.885 | 0.882 | 0.859 | 0.880 |
| | | IM | 0.927 | 1.0 | 0.995 | 0.962 | 0.938 | 0.966 | 0.941 | 0.996 | 0.947 | 0.966 |
| | | OM | 0.994 | 1.0 | 0.995 | 0.997 | 0.664 | 0.793 | 0.829 | 0.767 | 0.772 | 0.793 |
| **1024-YOLO26-LD** | | All | 0.992 | 1.0 | 0.995 | 0.996 | 0.868 | 0.925 | 0.918 | 0.937 | 0.912 | 0.926 |
| | | IM | 0.992 | 1.0 | 0.995 | 0.996 | 0.948 | 0.971 | 0.952 | 0.996 | 0.952 | 0.971 |
| | | OM | 0.992 | 1.0 | 0.995 | 0.996 | 0.788 | 0.880 | 0.884 | 0.878 | 0.872 | 0.880 |
| **640-U-Net-LD** | | All | 0.994 | 0.951 | 0.998 | 0.972 | 0.961 | 0.980 | 0.972 | 0.989 | 0.998 | 0.980 |
| | | IM | 0.992 | 0.944 | 0.997 | 0.967 | 0.956 | 0.977 | 0.968 | 0.988 | 0.997 | 0.977 |
| | | OM | 0.996 | 0.957 | 0.998 | 0.979 | 0.967 | 0.982 | 0.976 | 0.990 | 0.998 | 0.982 |
| **1024-U-Net-LD** | | All | 0.977 | 0.985 | 0.999 | 0.981 | 0.972 | 0.985 | 0.984 | 0.987 | 0.999 | 0.985 |
| | | IM | 0.975 | 0.981 | 0.999 | 0.970 | 0.966 | 0.983 | 0.983 | 0.983 | 0.999 | 0.983 |
| | | OM | 0.981 | 0.988 | 0.999 | 0.984 | 0.977 | 0.988 | 0.986 | 0.991 | 0.999 | 0.988 |
| **640-Detectron2-LD** | | All | 0.908 | 0.984 | 0.950 | 0.944 | 0.830 | 0.880 | 0.844 | 0.983 | 0.960 | 0.880 |
| | | IM | 0.903 | 0.981 | 0.934 | 0.940 | 0.828 | 0.881 | 0.844 | 0.980 | 0.938 | 0.881 |
| | | OM | 0.989 | 0.987 | 0.977 | 0.988 | 0.832 | 0.879 | 0.844 | 0.986 | 0.982 | 0.879 |
| **1024-Detectron2-LD** | | All | 0.945 | 0.943 | 0.961 | 0.944 | 0.589 | 0.684 | 0.599 | 0.985 | 0.959 | 0.684 |
| | | IM | 0.893 | 0.903 | 0.937 | 0.898 | 0.618 | 0.713 | 0.627 | 0.985 | 0.713 | 0.940 |
| | | OM | 0.993 | 0.978 | 0.975 | 0.985 | 0.560 | 0.655 | 0.571 | 0.985 | 0.655 | 0.978 |
| **640-SAM3-LD** | | All | 0.995 | 0.991 | 1.0 | 0.993 | 0.953 | 0.974 | 0.961 | 0.991 | 0.994 | 0.974 |
| | | IM | 0.992 | 0.990 | 1.0 | 0.991 | 0.950 | 0.972 | 0.958 | 0.991 | 0.993 | 0.972 |
| | | OM | 0.998 | 0.992 | 1.0 | 0.995 | 0.957 | 0.976 | 0.964 | 0.992 | 0.995 | 0.976 |
| **1024-SAM3-LD** | | All | 0.953 | 0.992 | 1.0 | 0.972 | 0.955 | 0.975 | 0.961 | 0.993 | 0.994 | 0.975 |
| | | IM | 0.994 | 0.991 | 1.0 | 0.993 | 0.951 | 0.973 | 0.959 | 0.992 | 0.993 | 0.973 |
| | | OM | 0.998 | 0.993 | 1.0 | 0.995 | 0.958 | 0.977 | 0.964 | 0.993 | 0.996 | 0.977 |

*All instance-based model outputs (YOLO, Detectron2 and SAM3) were calculated at IOU@0.5 with a swept confidence threshold at instance-level evaluation for mAP50. U-Net metrics are calculated at the best confidence with a swept confidence threshold with AUPRC, not mAP50 (also referred as mAP at IOU@0.5, instance metric only) at pixel-level evaluation. All metrics are globally pooled.
†All model outputs were converted to pixel-level and fixed at 0.5 confidence threshold (except for AUPRC sweeping across all confidence thresholds). All metrics are macro averaged per image.

**Supplementary Table 2.** Performance metrics on the valid images for the ultralow-dose (ULD) OM-IM fine-tuned models.

| Model | Electron Dose | Class | Mask Precision* | Mask Recall* | mAP50/ AUPRC* | Best F1-Score* | Mean IOU† | Mean Dice† | Mean Mask Precision† | Mean Mask Recall† | AUPRC† | Mean F1-Score† |
|---|---|---|---|---|---|---|---|---|---|---|---|---|
| **640-YOLOv11-ULD** | ULD | All | 0.952 | 0.952 | 0.947 | 0.952 | 0.831 | 0.902 | 0.907 | 0.897 | 0.900 | 0.902 |
| | | IM | 1.0 | 0.995 | 0.995 | 0.997 | 0.965 | 0.982 | 0.976 | 0.988 | 0.983 | 0.983 |
| | | OM | 0.904 | 0.909 | 0.898 | 0.906 | 0.698 | 0.821 | 0.838 | 0.805 | 0.817 | 0.821 |
| **1024-YOLOv11-ULD** | | All | 0.997 | 1.0 | 0.995 | 0.998 | 0.866 | 0.924 | 0.932 | 0.917 | 0.924 | 0.924 |
| | | IM | 0.998 | 1.0 | 0.995 | 0.999 | 0.973 | 0.986 | 0.985 | 0.988 | 0.987 | 0.986 |
| | | OM | 0.995 | 1.0 | 0.995 | 0.998 | 0.759 | 0.862 | 0.880 | 0.846 | 0.861 | 0.862 |
| **640-YOLO26-ULD** | | All | 0.934 | 0.949 | 0.986 | 0.941 | 0.821 | 0.894 | 0.889 | 0.902 | 0.892 | 0.894 |
| | | IM | 1.0 | 0.999 | 0.995 | 1.0 | 0.963 | 0.981 | 0.972 | 0.990 | 0.982 | 0.981 |
| | | OM | 0.868 | 0.899 | 0.977 | 0.883 | 0.678 | 0.808 | 0.805 | 0.814 | 0.802 | 0.808 |
| **1024-YOLO26-ULD** | | All | 1.0 | 0.954 | 0.991 | 0.976 | 0.824 | 0.898 | 0.927 | 0.874 | 0.900 | 0.898 |
| | | IM | 1.0 | 1.0 | 0.995 | 1.0 | 0.931 | 0.963 | 0.980 | 0.948 | 0.968 | 0.963 |
| | | OM | 1.0 | 0.907 | 0.988 | 0.951 | 0.717 | 0.834 | 0.873 | 0.801 | 0.832 | 0.834 |
| **640-U-Net-ULD** | | All | 0.982 | 0.977 | 0.997 | 0.980 | 0.961 | 0.980 | 0.979 | 0.982 | 0.997 | 0.980 |
| | | IM | 0.979 | 0.975 | 0.997 | 0.977 | 0.955 | 0.977 | 0.974 | 0.979 | 0.997 | 0.977 |
| | | OM | 0.986 | 0.978 | 0.998 | 0.982 | 0.967 | 0.983 | 0.982 | 0.981 | 0.998 | 0.983 |
| **1024-U-Net-ULD** | | All | 0.985 | 0.983 | 0.999 | 0.984 | 0.964 | 0.982 | 0.989 | 0.975 | 0.999 | 0.982 |
| | | IM | 0.980 | 0.980 | 0.998 | 0.980 | 0.957 | 0.978 | 0.985 | 0.971 | 0.999 | 0.978 |
| | | OM | 0.989 | 0.985 | 0.999 | 0.987 | 0.972 | 0.986 | 0.992 | 0.979 | 0.999 | 0.986 |
| **640-Detectron2-ULD** | | All | 0.976 | 0.972 | 0.957 | 0.974 | 0.653 | 0.782 | 0.664 | 0.977 | 0.961 | 0.782 |
| | | IM | 0.964 | 0.972 | 0.943 | 0.968 | 0.687 | 0.805 | 0.698 | 0.978 | 0.949 | 0.805 |
| | | OM | 0.987 | 0.972 | 0.968 | 0.979 | 0.619 | 0.758 | 0.629 | 0.976 | 0.973 | 0.758 |
| **1024-Detectron2-ULD** | | All | 0.985 | 0.950 | 0.956 | 0.967 | 0.433 | 0.601 | 0.439 | 0.972 | 0.960 | 0.601 |
| | | IM | 0.981 | 0.955 | 0.950 | 0.968 | 0.472 | 0.639 | 0.479 | 0.970 | 0.955 | 0.639 |
| | | OM | 0.989 | 0.959 | 0.962 | 0.974 | 0.955 | 0.563 | 0.399 | 0.973 | 0.965 | 0.563 |
| **640-SAM3-ULD** | | All | 0.994 | 0.984 | 1.0 | 0.989 | 0.978 | 0.989 | 0.994 | 0.983 | 0.99 | 0.989 |
| | | IM | 0.995 | 0.977 | 1.0 | 0.986 | 0.971 | 0.985 | 0.995 | 0.976 | 0.987 | 0.985 |
| | | OM | 0.994 | 0.991 | 1.0 | 0.993 | 0.985 | 0.992 | 0.994 | 0.991 | 0.993 | 0.992 |
| **1024-SAM3-ULD** | | All | 0.995 | 0.986 | 1.0 | 0.990 | 0.980 | 0.990 | 0.995 | 0.985 | 0.990 | 0.990 |
| | | IM | 0.995 | 0.979 | 1.0 | 0.987 | 0.974 | 0.987 | 0.995 | 0.978 | 0.987 | 0.987 |
| | | OM | 0.995 | 0.991 | 1.0 | 0.993 | 0.986 | 0.993 | 0.995 | 0.991 | 0.994 | 0.993 |

*All instance-based model outputs (YOLO, Detectron2 and SAM3) were calculated at IOU@0.5 with a swept confidence threshold at instance-level evaluation for mAP50. U-Net metrics are calculated at the best confidence with a swept confidence threshold with AUPRC, not mAP50 (also referred as mAP at IOU@0.5, instance metric only) at pixel-level evaluation. All metrics are globally pooled.

†All model outputs were converted to pixel-level and fixed at 0.5 confidence threshold (except for AUPRC sweeping across all confidence thresholds). All metrics are macro averaged per image.

**Supplementary Table 3.** Top 10 models in the ULD seed sorted by best F1-score from the Top Model Decision Tree. Both low-dose and ultralow-dose test images were inferenced for a top 10 model screen. F1-scores are colored by gradient (green at ≥ 0.75, yellow at ≥ 0.5, red at ≥ 0.25 and black at 0).

| Model | Model Image Size | Test Image Size | Model Electron Dose | Test Electron Dose | Class | Mask Level | Mask Precision* | Mask Recall* | mAP50/ AUPRC* | Best F1-Score* | Mean IOU† | Mean Dice† | Mean Mask Precision† | Mean Mask Recall† | AUPRC† | Mean F1-Score† | FPS |
|---|---|---|---|---|---|---|---|---|---|---|---|---|---|---|---|---|---|
| 1024-YOLOv11 | 1024 | 1024 | LD | LD | All | object | 0.997 | 1.000 | 0.995 | 0.999 | 0.877 | 0.930 | 0.930 | 0.931 | 0.935 | 0.930 | 47 |
| 640-YOLOv11 | 640 | 1024 | LD | LD | All | object | 0.996 | 1.000 | 0.995 | 0.998 | 0.888 | 0.937 | 0.934 | 0.941 | 0.938 | 0.937 | 46 |
| 1024-SAM3 | 1024 | 1024 | LD | LD | All | object | 0.995 | 0.993 | 1.000 | 0.994 | 0.990 | 0.995 | 0.996 | 0.994 | 0.994 | 0.995 | 8 |
| 640-SAM3 | 640 | 1024 | LD | LD | All | object | 0.994 | 0.993 | 1.000 | 0.994 | 0.989 | 0.994 | 0.996 | 0.993 | 0.995 | 0.994 | 8 |
| 1024-U-Net | 1024 | 1024 | LD | LD | All | pixel | 0.985 | 0.990 | 0.997 | 0.987 | 0.972 | 0.985 | 0.990 | 0.982 | 0.998 | 0.985 | 58 |
| 640-U-Net | 640 | 1024 | LD | LD | All | pixel | 0.979 | 0.984 | 0.996 | 0.982 | 0.966 | 0.983 | 0.975 | 0.990 | 0.995 | 0.983 | 58 |
| 640-Detectron2 | 640 | 1024 | LD | LD | All | object | 0.973 | 0.988 | 0.967 | 0.980 | 0.957 | 0.978 | 0.972 | 0.985 | 0.971 | 0.978 | 33 |
| 1024-Detectron2 | 1024 | 1024 | LD | LD | All | object | 0.935 | 0.981 | 0.965 | 0.957 | 0.626 | 0.732 | 0.633 | 0.988 | 0.971 | 0.732 | 33 |
| 1024-YOLO26 | 1024 | 1024 | LD | LD | All | object | 0.921 | 0.975 | 0.966 | 0.947 | 0.876 | 0.929 | 0.926 | 0.935 | 0.931 | 0.929 | 41 |
| 640-YOLO26 | 640 | 1024 | LD | LD | All | object | 0.775 | 0.800 | 0.814 | 0.787 | 0.700 | 0.782 | 0.875 | 0.734 | 0.825 | 0.782 | 42 |
| 640-SAM3 | 640 | 1024 | LD | ULD | All | object | 0.821 | 0.916 | 0.752 | 0.866 | 0.761 | 0.840 | 0.913 | 0.842 | 0.913 | 0.840 | 8 |
| 1024-SAM3 | 1024 | 1024 | LD | ULD | All | object | 0.850 | 0.877 | 0.728 | 0.863 | 0.760 | 0.835 | 0.891 | 0.863 | 0.910 | 0.835 | 8 |
| 1024-U-Net | 1024 | 1024 | LD | ULD | All | pixel | 0.657 | 0.680 | 0.649 | 0.669 | 0.484 | 0.594 | 0.614 | 0.619 | 0.650 | 0.594 | 58 |
| 1024-Detectron2 | 1024 | 1024 | LD | ULD | All | object | 0.654 | 0.389 | 0.459 | 0.488 | 0.223 | 0.350 | 0.238 | 0.832 | 0.473 | 0.350 | 32 |
| 640-U-Net | 640 | 1024 | LD | ULD | All | pixel | 0.248 | 0.402 | 0.274 | 0.307 | 0.025 | 0.043 | 0.210 | 0.026 | 0.280 | 0.043 | 58 |
| 640-Detectron2 | 640 | 1024 | LD | ULD | All | object | 0.934 | 0.179 | 0.295 | 0.300 | 0.448 | 0.531 | 0.629 | 0.496 | 0.614 | 0.531 | 32 |
| 1024-YOLOv11 | 1024 | 1024 | LD | ULD | All | object | 0.838 | 0.161 | 0.243 | 0.270 | 0.191 | 0.236 | 0.315 | 0.206 | 0.553 | 0.236 | 47 |
| 1024-YOLO26 | 1024 | 1024 | LD | ULD | All | object | 0.227 | 0.200 | 0.257 | 0.212 | 0.107 | 0.134 | 0.228 | 0.112 | 0.540 | 0.134 | 43 |
| 640-YOLOv11 | 640 | 1024 | LD | ULD | All | object | 0.400 | 0.036 | 0.222 | 0.067 | 0.029 | 0.035 | 0.075 | 0.032 | 0.500 | 0.035 | 48 |
| 640-YOLO26 | 640 | 1024 | LD | ULD | All | object | 0.000 | 0.000 | 0.000 | 0.000 | 0.000 | 0.000 | 0.000 | 0.000 | 0.041 | 0.000 | 45 |

*All instance-based model outputs (YOLO, Detectron2 and SAM3) were calculated at IOU@0.5 with a swept confidence threshold at instance-level evaluation for mAP50. U-Net metrics are calculated at the best confidence with a swept confidence threshold with AUPRC, not mAP50 (also referred as mAP at IOU@0.5, instance metric only) at pixel-level evaluation. All metrics are globally pooled.

†All model outputs were converted to pixel-level and fixed at 0.5 confidence threshold (except for AUPRC sweeping across all confidence thresholds).All metrics are macro averaged per image.

**Supplementary Table 4.** Top 10 models in the ULD seed sorted by best F1-score from the Top Model Decision Tree. Both low-dose and ultralow-dose test images were inferenced for a top 10 model screen. F1-scores are colored by gradient (green at ≥ 0.75, yellow at ≥ 0.5, red at ≥ 0.25 and black at 0).

| Model | Model Image Size | Test Image Size | Model Electron Dose | Test Electron Dose | Class | Mask Level | Mask Precision* | Mask Recall* | mAP50/ AUPRC* | Best F1-Score* | Mean IOU† | Mean Dice† | Mean Mask Precision† | Mean Mask Recall† | AUPRC† | Mean F1-Score† | FPS |
|---|---|---|---|---|---|---|---|---|---|---|---|---|---|---|---|---|---|
| 1024-SAM3 | 1024 | 640 | ULD | LD | All | object | 0.992 | 0.981 | 1.000 | 0.986 | 0.973 | 0.986 | 0.994 | 0.980 | 0.984 | 0.986 | 8 |
| 640-SAM3 | 640 | 640 | ULD | LD | All | object | 0.992 | 0.979 | 1.000 | 0.986 | 0.972 | 0.986 | 0.994 | 0.978 | 0.986 | 0.986 | 8 |
| 640-Detectron2 | 640 | 640 | ULD | LD | All | object | 0.983 | 0.962 | 0.961 | 0.972 | 0.706 | 0.801 | 0.721 | 0.977 | 0.968 | 0.801 | 34 |
| 640-U-Net | 640 | 640 | ULD | LD | All | pixel | 0.956 | 0.940 | 0.982 | 0.948 | 0.893 | 0.936 | 0.975 | 0.913 | 0.981 | 0.936 | 135 |
| 1024-Detectron2 | 1024 | 640 | ULD | LD | All | object | 0.923 | 0.969 | 0.950 | 0.946 | 0.535 | 0.671 | 0.543 | 0.981 | 0.956 | 0.671 | 35 |
| 1024-U-Net | 1024 | 640 | ULD | LD | All | pixel | 0.860 | 0.887 | 0.957 | 0.873 | 0.776 | 0.840 | 0.975 | 0.792 | 0.955 | 0.840 | 129 |
| 1024-YOLOv11 | 1024 | 640 | ULD | LD | All | object | 0.868 | 0.875 | 0.856 | 0.872 | 0.762 | 0.848 | 0.873 | 0.837 | 0.841 | 0.848 | 64 |
| 1024-YOLO26 | 1024 | 640 | ULD | LD | All | object | 0.857 | 0.844 | 0.828 | 0.851 | 0.756 | 0.843 | 0.858 | 0.839 | 0.838 | 0.843 | 58 |
| 640-YOLO26 | 640 | 640 | ULD | LD | All | object | 0.784 | 0.788 | 0.731 | 0.786 | 0.675 | 0.768 | 0.813 | 0.746 | 0.785 | 0.768 | 58 |
| 640-YOLOv11 | 640 | 640 | ULD | LD | All | object | 0.749 | 0.748 | 0.637 | 0.749 | 0.710 | 0.801 | 0.850 | 0.772 | 0.799 | 0.801 | 65 |
| 1024-SAM3 | 1024 | 640 | ULD | ULD | All | object | 0.901 | 0.911 | 0.897 | 0.906 | 0.810 | 0.877 | 0.915 | 0.885 | 0.935 | 0.877 | 8 |
| 640-SAM3 | 640 | 640 | ULD | ULD | All | object | 0.875 | 0.916 | 0.881 | 0.895 | 0.803 | 0.874 | 0.889 | 0.905 | 0.925 | 0.874 | 8 |
| 640-Detectron2 | 640 | 640 | ULD | ULD | All | object | 0.836 | 0.744 | 0.790 | 0.787 | 0.543 | 0.684 | 0.597 | 0.873 | 0.820 | 0.684 | 30 |
| 1024-Detectron2 | 1024 | 640 | ULD | ULD | All | object | 0.662 | 0.782 | 0.679 | 0.717 | 0.419 | 0.561 | 0.449 | 0.844 | 0.691 | 0.561 | 35 |
| 1024-U-Net | 1024 | 640 | ULD | ULD | All | pixel | 0.635 | 0.781 | 0.737 | 0.701 | 0.319 | 0.431 | 0.811 | 0.372 | 0.734 | 0.431 | 138 |
| 1024-YOLOv11 | 1024 | 640 | ULD | ULD | All | object | 0.718 | 0.645 | 0.668 | 0.680 | 0.600 | 0.718 | 0.772 | 0.716 | 0.750 | 0.718 | 64 |
| 1024-YOLO26 | 1024 | 640 | ULD | ULD | All | object | 0.652 | 0.542 | 0.562 | 0.591 | 0.534 | 0.655 | 0.754 | 0.629 | 0.702 | 0.655 | 55 |
| 640-YOLOv11 | 640 | 640 | ULD | ULD | All | object | 0.784 | 0.464 | 0.559 | 0.583 | 0.447 | 0.572 | 0.764 | 0.489 | 0.678 | 0.572 | 63 |
| 640-U-Net | 640 | 640 | ULD | ULD | All | pixel | 0.617 | 0.397 | 0.460 | 0.483 | 0.151 | 0.205 | 0.590 | 0.161 | 0.459 | 0.205 | 134 |
| 640-YOLO26 | 640 | 640 | ULD | ULD | All | object | 0.594 | 0.399 | 0.451 | 0.477 | 0.335 | 0.427 | 0.586 | 0.388 | 0.613 | 0.427 | 58 |

*All instance-based model outputs (YOLO, Detectron2 and SAM3) were calculated at IOU@0.5 with a swept confidence threshold at instance-level evaluation for mAP50. U-Net metrics are calculated at the best confidence with a swept confidence threshold with AUPRC, not mAP50 (also referred as mAP at IOU@0.5, instance metric only) at pixel-level evaluation. All metrics are globally pooled.

†All model outputs were converted to pixel-level and fixed at 0.5 confidence threshold (except for AUPRC sweeping across all confidence thresholds).All metrics are macro averaged per image.

### References

Madugula, S. S., Massenburg, L. N., Brown, S. R., Bible, A. N., Harris, C. R., Zhang, L. X., Parker, K., Retterer, S. T., Morrell-Falvey, J. L., Vasudevan, R. K., & Williams, A. N. (2026). Automated Bacterial Identification and Morphological Feature Analysis in Low-Dose Cryo-EM Using YOLOv11. *Advanced Intelligent Discovery*, *n/a*(n/a), e202500241. https://doi.org/https://doi.org/10.1002/aidi.202500241
